# AF-CALVADOS: AlphaFold-guided simulations of multi-domain proteins at the proteome level

**DOI:** 10.1101/2025.10.19.683306

**Authors:** Sören von Bülow, Kristoffer E. Johansson, Kresten Lindorff-Larsen

**Affiliations:** Structural Biology and NMR Laboratory, Linderstrøm-Lang Centre for Protein Science, Department of Biology, University of Copenhagen, Copenhagen, Denmark

## Abstract

Deep-learning methods have transformed our ability to predict the three-dimensional structures of folded proteins from sequence, and coarse-grained simulations have made it possible to study intrinsically disordered proteins at the proteome scale. More than half of human proteins, however, contain mixtures of disordered regions and one or more folded domains, and the biological function of these multi-domain proteins depends on the interplay between the folded and disordered regions. Here, we developed AF-CALVADOS, a coarse-grained simulation model that is informed by AlphaFold to model the dynamics of intrinsically disordered proteins and multi-domain proteins containing mixtures of folded and disordered regions. AF-CALVADOS leverages information from AlphaFold 2 to model folded regions that we then integrate with the coarse-grained CALVADOS model. Our automated framework makes it possible to perform simulations of any soluble folded or disordered protein without manually defining the folded regions, enabling scaling to the proteome level. We validate AF-CALVADOS using experimental SAXS data for more than 400 proteins and find that it performs well across proteins with varying amounts of ordered and disordered regions. We demonstrate the scalability of AF-CALVADOS by performing simulations of 12,483 intracellular human proteins and make the data freely available; we envisage that large-scale simulation data generated by AF-CALVADOS can be used to benchmark or train machine learning models for flexible, multi-domain proteins. The conformational ensembles can also be used to study sequence-dynamics-function relationships at scale, and can shed light on the interplay between folded and disordered regions. We exemplify this by analysing the disordered regions in 1,487 human transcription factors.

## 1 Introduction

Structure prediction methods such as AlphaFold 2 (AF2) have made it possible to predict the structures of folded domains with close to experimental accuracy, and can be applied at the proteome scale^1^. These methods are complemented by powerful simulation or machine learning models that can be used to generate conformational ensembles of disordered regions at high speed and accuracy^2^. Most proteins, however, consist of mixtures of disordered regions and folded domains^3,4^, and neither AF2 nor most models for disordered regions capture such proteins.

Among the different possible coarse-grained simulation models, hydrophobicity scale (HPS) models have been used extensively for coarse-grained implicit solvent simulations of intrinsically disordered proteins and regions (IDPs and IDRs)^5–7^. These models map the protein structure to one amino-acid-specific spherical bead per residue. The 20 different beads are characterized by their size (*σ*), charge (*q*), and a *λ* value, which together capture contributions from van-der-Waals interactions, hydrogen bonds, *π*-*π* interactions and charge interactions. These models strike a balance of accuracy and speed, and have been used to perform proteome-level simulations of disordered proteins ^8^.

CALVADOS is one such HPS model that has been systematically parameterised against experimental small-angle X-ray scattering (SAXS) and nuclear magnetic resonance (NMR) data to predict single-chain global properties (radius of gyration, end-to-end distance) of a diverse set of IDRs^7^. Further, multi-chain simulations with the updated CALVADOS 2 model accurately predict phase coexistence properties (saturation concentrations) of IDRs ^9^. CALVADOS and other related HPS models, as well as most other coarse-grained models, cannot by default represent protein folded domains accurately. This is because the stability of the folded domains depends sensitively on the interactions of all atoms within the folded domains and between folded domains and solvent. Therefore, folded domains are generally unstable in coarse-grained simulations with HPS models. CALVADOS 3 is an extension of CALVADOS that enables the simulation of multi-domain proteins (MDPs) by restraining the folded domains with a harmonic elastic network and representing residues in the folded domains by their centres of mass^10^. The CALVADOS 3 force field was re-optimised to match single-chain experimental properties from SAXS and paramagnetic relaxation enhancement (PRE) NMR spectroscopy, and was found in the recent CASP16 to perform well in a blind assessment of the global conformational properties of two related MDPs ^11^. A key step in developing CALVADOS 3 was to represent residues in folded domains using the centre-of-mass positions of their constituent atoms. Despite these advances, it remains difficult to use HPS models for large-scale applications to generate conformational ensembles for thousands of MDPs. In particular, in previous work with CALVADOS 3 we used manual definitions of which residues of the proteins constitute folded domains and should be restrained. Furthermore, the exact definition of flexible linkers within a protein depends on somewhat subjective choices.

Beyond AF2’s accuracy in predicting the structures of folded domains, the predicted local distance difference (pLDDT) metric, designed to predict the local accuracy of the structure prediction, is an excellent predictor for the extent of local order vs. disorder ^12^. This, for example, enabled us to use AF2 pLDDT scores to define disordered regions of the entire human proteome and subsequently simulate them with CALVADOS 2^8^.

The AF2 predicted aligned error (PAE) score is a prediction of the alignment error of pairs of residues in a protein. Beyond the use for assessing the accuracy of structure prediction, it has been shown that the PAE scores correlate with fluctuations observed in molecular dynamics simulations^8,13,14^. Here, low PAE values indicate a relatively fixed distance between the residues, for example because they are part of the same domain.

The relationship between structure, disorder, fluctuations and the AF2 pLDDT and PAE scores have been used to study protein dynamics. For example, the pLDDT scores and PAE scores have been used in a method called AF-ENM to define protein-specific harmonic restraints in simulations with the coarse-grained MARTINI 3 forcefield^14^. Similarly, AlphaFold-Metainference uses data from AF2 together with CALVADOS to generate conformational ensembles of flexible proteins^15^, and Ensemblify can combine AF2 structures and PAE scores in Rosetta-based sampling^16^. AFflecto uses pLDDT scores to define disordered regions to be sampled^17^, and pLDDT scores have also been used to set native-bias constraints in CABS-flex simulations^18^. A Bayesian framework—called bAIes—combines AF2 predictions (distograms) with molecular simulations, and was shown to provide an accurate description of both local and global structure of disordered proteins and MDPs^19^. Finally, certain deep-learning architectures such as BioEMU^20^ and PepTron^21^ are now beginning to be able to model proteins with mixtures of folded and disordered regions, and models such as IDPForge can generate conformational ensembles of MDPs with manual^22^ or automated^23^ specification of the folded regions.

Extending these ideas, we here present AF-CALVADOS that combines AF2-based restraining of folded domains with the accuracy and speed of the CALVADOS models. Our goal with developing AF-CALVADOS was to create an accurate, robust, computationally-efficient and scalable approach of generating conformational ensembles of both disordered and MDPs. We validate AF-CALVADOS on a wide range of MDPs and show that it captures conformational ensembles in excellent agreement with experiments including the recently described PeptoneDB-SAXS benchmark^21^. We exploit the speed and accuracy of AF-CALVADOS to generate conformational ensembles of 12,483 intracellular human proteins with both folded and disordered regions, and analyse in more detail the ensembles of 1,487 human transcription factors.

## Results and Discussion

### Considerations when developing AF-CALVADOS

CALVADOS 3 was developed and parameterised to simulate IDRs and MDPs. For the simulation of MDPs, harmonic restraints are applied in CALVADOS 3 between close (*<* 0.9 nm) non-bonded residues in folded domains. The folded domains are defined manually and the restraining force constant is global and fixed, i.e., residue pairs are either fully restrained or not restrained.

We aimed to extend CALVADOS 3 with an automated restraining protocol using information from AF2. AF-CALVADOS applies restraints with gradual strength based on the AF2 pLDDT and PAE values. The pLDDT value denotes the residue-level confidence of the atomic positions of AF2, but is also correlated with local disorder. A high pLDDT value thus indicates that a residue forms substantial secondary and tertiary structure (possibly induced via intermolecular interactions). The PAE indicates the positional error of two residues relative to one another, but also correlates with the formation of structure and level of dynamics^14^. AF-CALVADOS thus extracts information from the AF2 pLDDT and PAE values to define the conformational restraints on folded domains rather than defining binary domain boundaries manually (Fig. 1). To enable a wider range of fluctuations, we use a 12-10 Gō-type potential for the structure-based restraints instead of the harmonic restraints used in CALVADOS 3.

**Figure 1:**
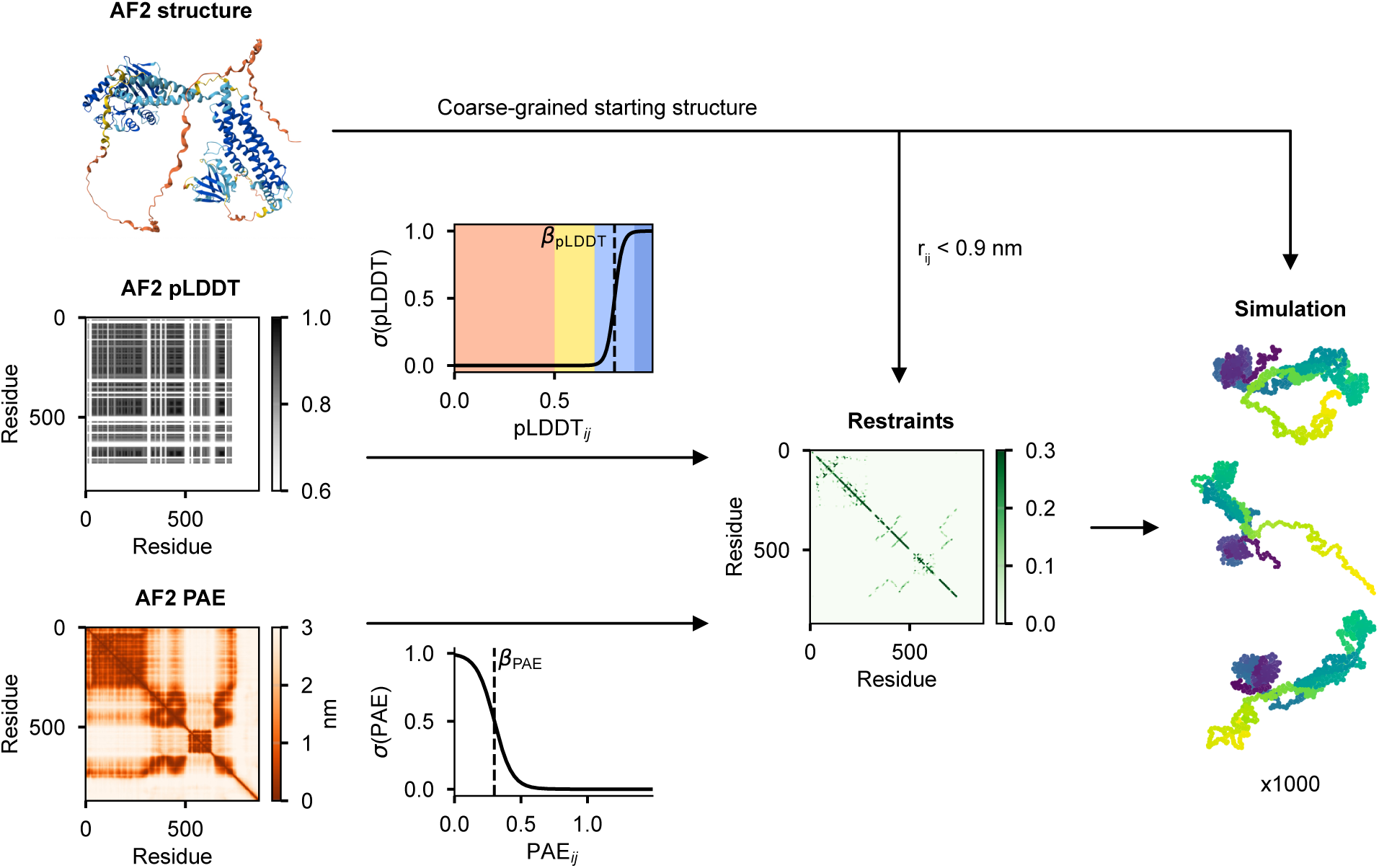
AF-CALVADOS restraining and simulation protocol based on AF2 structure, PAE and pLDDT. The AF2 structure is coarse-grained into a COM representation. Residue pairs with COM distance *<* 0.9 nm are considered for restraining. The AF2 pLDDT and PAE data are combined into restraints via switching functions *σ*. The AF-CALVADOS simulation records 1000 largely uncorrelated protein configurations. The colour scale shown on the pLDDT plot follows the AF2 convention with red (very low confidence, pLDDT*/*100 *<* 0.5), yellow (low confidence, 0.5 *<*pLDDT*/*100 *<* 0.5), light blue (high confidence, 0.7 *<*pLDDT*/*100 *<* 0.9) and dark blue (very high confidence pLDDT*/*100 *>* 0.9).

We used the AF2 pLDDT and PAE values in the restraining procedure (see Methods) based on the following considerations: (i) Unrestrained residues should behave as in CALVADOS 3. (ii) Only folded domains composed of residues predicted with high (0.7 *<*pLDDT*/*100 *<* 0.9) or very high confidence (pLDDT*/*100 *>* 0.9) should be restrained. The restraining potential should gradually increase in the ‘high’ confidence region and be maximal for pLDDT*/*100 *>* 0.9. (iii) Only pairs of residues with low PAE score should be restrained, as only those are predicted to undergo coordinated movements. We further use the PAE as a proxy for the fluctuation of distances in the residue pair in a conformational ensemble. (iv) For restrained pairs of residues we do not include the CALVADOS non-electrostatic and electrostatic interaction. Edge cases (very low restraining potential) are accounted for by interpolating between restraining potential and the CALVADOS non-bonded potentials. (v) Equilibrium bond distances of fully unrestrained residues are as in CALVADOS 3 (0.38 nm). Equilibrium bond distances in folded (restrained) regions are taken as the distances between the centres of mass in the starting structure generated for example with AF2. Edge cases are accounted for by interpolating between bond distances from CALVADOS 3 and those from the starting PDB structure, as in (iv). We used switching functions to convert pLDDT and PAE values into restraints, as well as treating edge cases. The parameters for the switching function for including pLDDT information were fixed to exclude non-confident structure predictions.

As described below, we optimised the parameters for the strength of the structure-based potential (*t:*), and the midpoint (*β*_PAE_) and steepness (*α*_PAE_) of the PAE switching function, using two criteria: (i) Simulations of MDPs should be in good agreement with experimentally determined values of the radius of gyration (*R*_g_). If too many residues are considered folded/unfolded, the calculated *R*_g_ should be strongly affected. (ii) The inter-residue fluctuations should reflect the PAE matrix from AF2. We therefore optimised the parameters such that domains with low PAE values also have low pair distance fluctuations in the simulations.

### Parameterising AF-CALVADOS using experimental data

We performed a grid scan of the parameters *t:* and *β*_PAE_ for two values of *α*_PAE_ (15 nm*^−^*^1^ and 30 nm*^−^*^1^). We show the (very similar) results corresponding to *α*_PAE_ = 15 nm*^−^*^1^ and 30 nm*^−^*^1^ in Fig. 2A,B and SI Fig. S1A,B, respectively. For each combination of parameters we performed simulations for a set of MDPs with experimental data^10^ and compared the simulation results to experiments. The results show the agreement quantified via a *χ*^2^ value and the absolute relative error, which considers both the magnitude and direction of the deviations between simulation and simulated *R*_g_ values. Overall, the automatic restraining procedure leads to excellent agreement between simulation and experimental *R*_g_ for a range of parameters (SI Fig. S2). The variation over our 1000-fold bootstrapping show that several parameter combinations lead to similar agreement between simulation and experiments. Combinations including *β*_PAE_ = 0.3 nm appear to strike a balance between low *χ*^2^ and a relative error close to zero.

**Figure 2:**
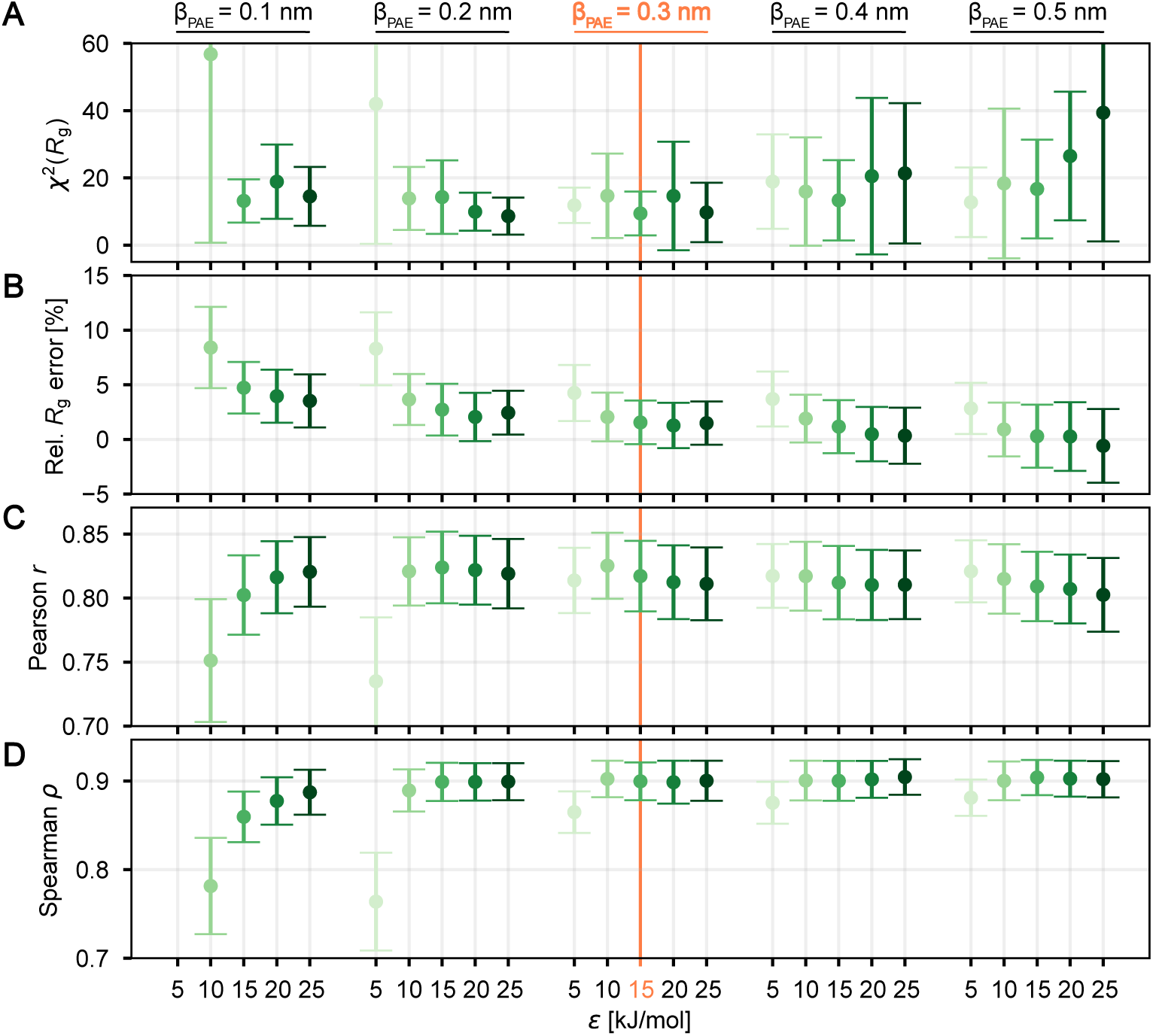
AF-CALVADOS parameterisation with *α*_PAE_ = 15 nm*^−^*^1^. (A, B) Agreement with experimental *R*_g_ data on 23 proteins10. (C, D) Agreement using metrics describing the recovery of AF2 PAE matrix for 30 diverse proteins. Error bars indicate 95% confidence intervals estimated by 1000-fold bootstrapping. The orange line indicates data generated with *β*_PAE_ = 0.3 nm and *t:* = 15 kJ mol*^−^*^1^, which is the parameter set that we selected from this analysis.

### Parameter selection to recover folded domains as described by the PAE

We applied the same parameter scan as above to 30 randomly selected proteins from a set of non-transmembrane intracellular proteins (see Methods) at physiological temperature (*T* = 310 K). We compared the matrix *σ*(*r*) of standard deviations of simulation distance fluctuations between all residues in the protein to the PAE map that was used to define the restraints.

As an example, we show a subset of the PAE and *σ*(*r*) matrices for the transcription elongation factor TCEA3 (Uniprot entry: O75764; Fig. 3) and the full parameter scan in SI Fig. S3. Too low values of *β*_PAE_ and *t:* lead to unfolding of one or both of the folded domains, but many parameter combinations lead to qualitatively similar agreement between the PAE and *σ*(*r*).

**Figure 3:**
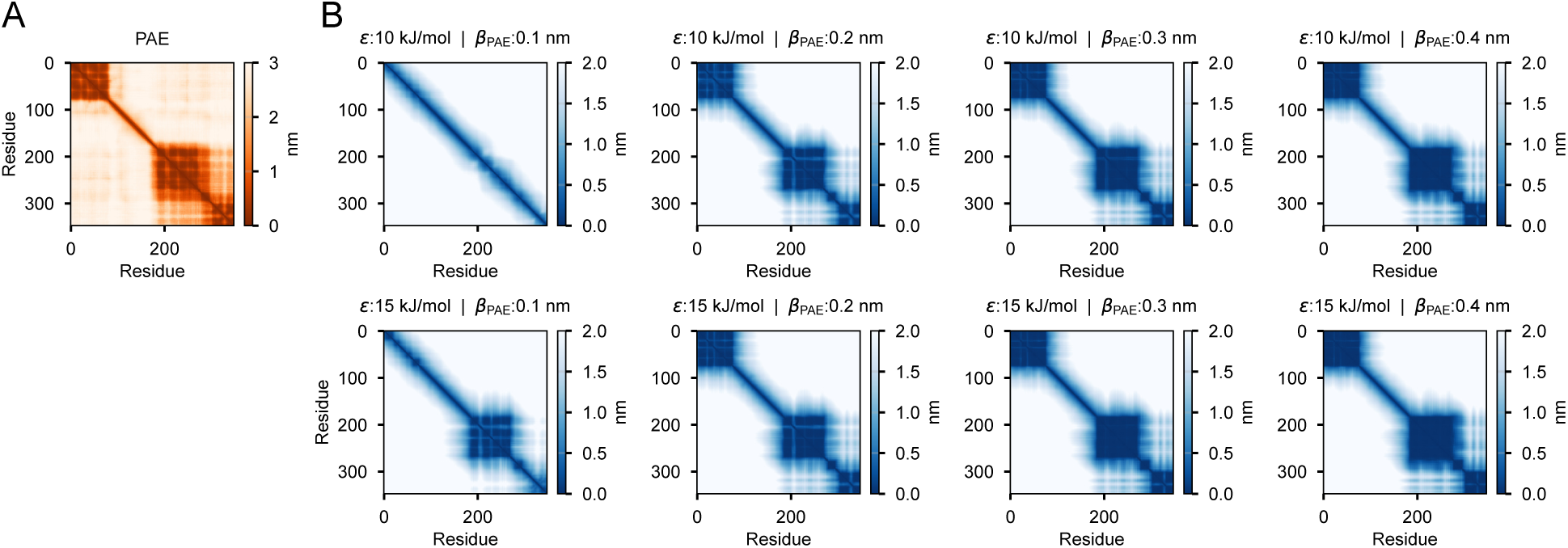
Comparison of the PAE matrix (A) vs. set of *σ*(*r*) matrices (B) illustrated by the transcription elongation factor TCEA3 (Uniprot entry: O75764). *t:* and *β*_PAE_ values indicate the parameters used in the simulation. Results are shown with *α*_PAE_ = 15 nm*^−^*^1^.

We note that we do not expect a perfect correlation between PAE and *σ*(*r*), as some correlations reported in the PAE are between poorly resolved low-confidence (low pLDDT) domains, which we intentionally do not restrain (see above consideration (ii)). Further, since PAE is a measure of prediction accuracy, it is only indirectly a probe of residue–residue interactions and fluctuations^8,13,14^. The results for the 30 proteins show that many parameter combinations, including those with *β*_PAE_ = 0.3 nm, show very good correlation between PAE and *σ*(*r*) values (Fig. 2C+D and SI Fig. S1C+D).

Based on the comparisons to the experimental *R*_g_ values and the AF2 PAE values (Fig. 2 and SI Fig. S1) we selected *α*_PAE_ = 15 nm*^−^*^1^, *β*_PAE_ = 0.3 nm, and *t:* = 15 kJ mol*^−^*^1^ as parameters for AF-CALVADOS. This parameter set performed well across all metrics reporting on agreement to experimental chain expansion and PAE values; however, several other parameter sets perform comparably to the chosen values, indicating robustness of the model.

### Benchmarking AF-CALVADOS

Having developed and parameterised an approach to combine AF2 and CALVADOS we assess the final model. We first evaluate the accuracy on capturing experimental chain expansion for 23 MDPs and compare with the original CALVADOS 3 model (with manual domain definitions). The results show that AF-CALVADOS simulations are in good agreement with experiments and perform as well as the CALVADOS 3 model (Fig. 4).

**Figure 4:**
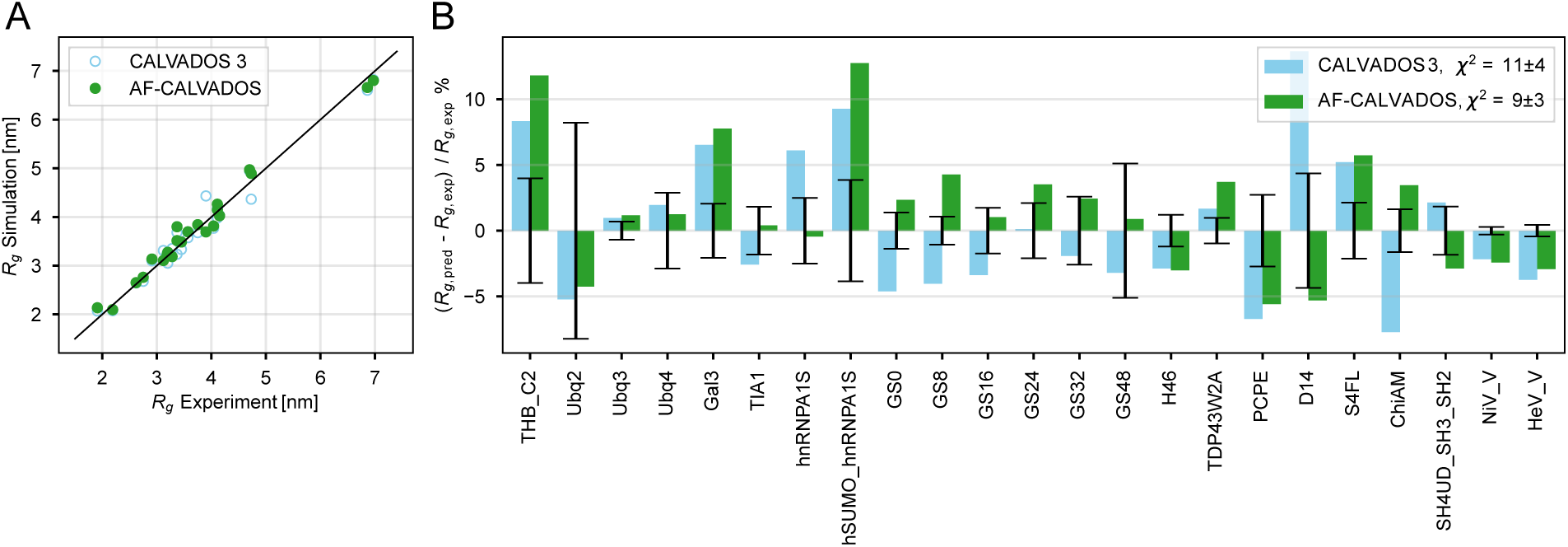
Agreement of AF-CALVADOS *R*_g_ (green) and CALVADOS 3 *R*_g_ (light blue)10 with experimental *R*_g_. (A) Scatter plot of *R*_g_ data. (B) Relative error between simulation and experimental *R*_g_. Black error bars indicate the reported experimental errors.

To test AF-CALVADOS more broadly and independently of the proteins used to develop it, we used the recently described PeptoneBench SAXS benchmark^21^. Specifically, the PeptoneDB-SAXS dataset consists of 439 proteins with different amounts of protein disorder^21^ and with SAXS data available in SASBDB^24^. We used AF-CALVADOS to generate conformational ensembles of these proteins, and compared the results (after removing proteins with high similarity to the CALVADOS training data; see Methods) to previously generated benchmark results^21^ for a series of generative models (Fig. 5). AF2^25^ and Boltz2^26^ were developed to predict individual structures for folded proteins and give rise to excellent agreement with SAXS data for proteins with little disorder, but show lower accuracy for proteins with substantial disorder (Fig. 5). Conversely, the IDP-o model was developed to predict structural ensembles of disordered proteins^21^ and shows the greatest accuracy for these proteins (Fig. 5). In contrast, BioEMU^20^ and PepTron^21^ provide similar accuracy (at the resolution afforded by the SAXS data) to AF2 for folded proteins, but substantially improved accuracy for more disordered proteins (Fig. 5). Finally, we find that AF-CALVADOS provides excellent accuracy across the entire order–disorder scale (Fig. 5). These observations can be quantified by the area under the LOWESS curve as a measure of the accuracy across disorder contents^21^ showing that AF-CALVADOS performs better than the other methods (Fig. 5). The accuracy can be improved further by using 1000 AF-CALVADOS structures instead of 100 (SI Fig. S4). A complementary application of the PeptoneDB-SAXS benchmark is to assess the accuracy of the ensembles after reweighting the ensembles with the experimental data^21^, and AF-CALVADOS also provides broad and high accuracy in this test (SI Figs. S4 and S5).

**Figure 5:**
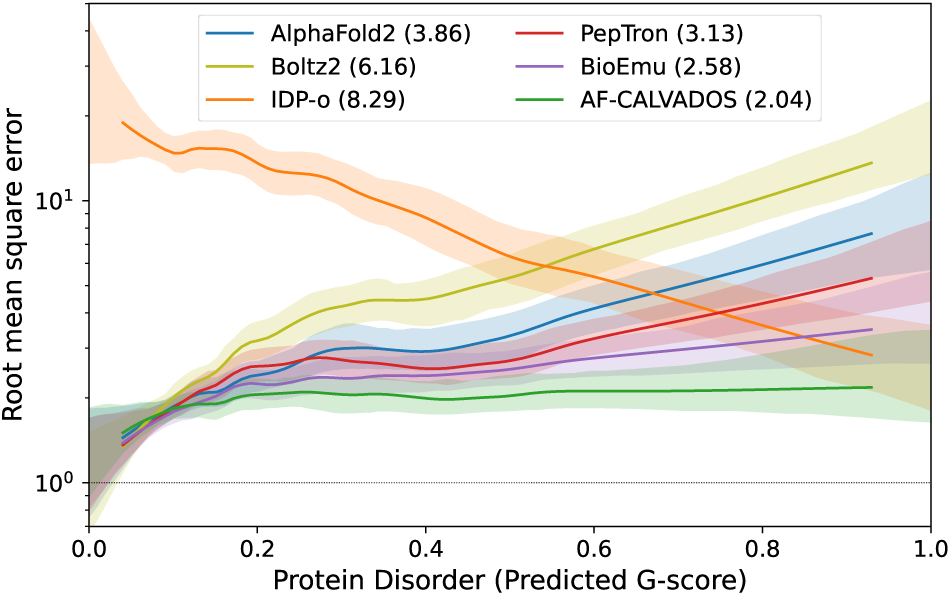
Benchmarking AF-CALVADOS using experimental SAXS data of 410 proteins. The figure shows a summary of the PeptoneBench SAXS benchmark where conformational ensembles are compared to measured SAXS data via back-calculated scattering intensities. Each line is a locally weighted scatterplot smoothed (LOWESS) curve that summarizes the root-mean-square-error of the calculated SAXS intensities across proteins with different levels of disorder. The latter is quantified by the predicted G-score that ranges from zero for fully ordered proteins to one for completely disordered proteins. The area under the LOWESS curve summarizes the accuracy across the entire order–disorder continuum21 and is provided in the legend. This figure shows the results for ensembles with 100 conformations, without filtering or reweighting, for 410 proteins that show ≤ 80% sequence identity to proteins used to parameterise CALVADOS (see Methods for details). Other results for the benchmark are shown in the Supporting Information.

Having compared AF-CALVADOS to independent experimental SAXS data, we then compared simulations with the final AF-CALVADOS model with the PAE and *σ*(*r*) matrices for the set of 30 intracellular proteins described above and for an additional set of 30 randomly selected transcription factors not used during parameterisation, and find that the AF-CALVADOS simulations capture the structural features defined by the PAE matrices (SI Figs. S6 and S7). Finally, we tested the agreement between the PAE and *σ*(*r*) from AF-CALVADOS simulations of three proteins with Zinc-finger domains (Fig. 6). We consider these domains as a particular challenging test case due to their small size and their stabilisation by zinc ions not explicitly represented by AF2 or CALVADOS. We find that the Zinc-finger domains in the Oxysterol receptor LXR-beta (P55055) and the Insulin gene enhancer (P61371) are recovered well; however, only 12 of the 15 Zinc-finger domains in REPIN1 (Q9BWE0) are stable in simulations because the three most N-terminal domains have low AF2 pLDDT values. In summary, if AF2 predicts Zinc-fingers domains with high confidence, AF-CALVADOS maintains these domains in the simulation, but if AF2 has low confidence predictions the domains will be unstable in the simulations.

**Figure 6:**
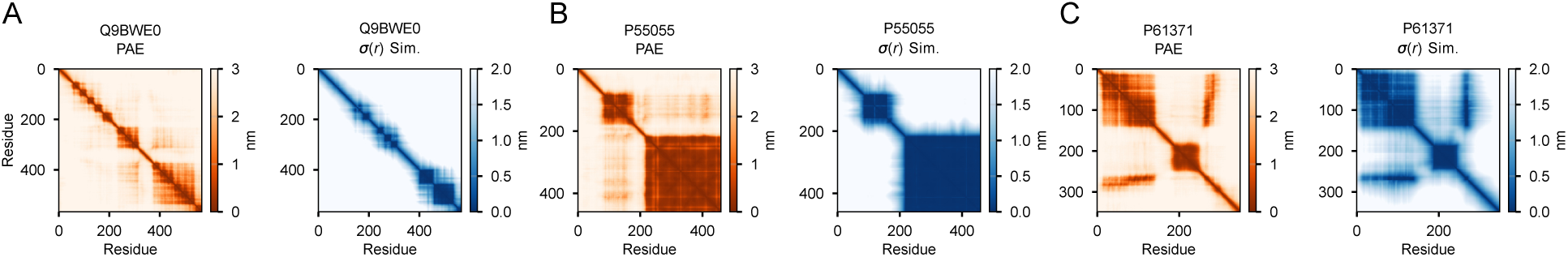
PAE matrix (orange) and *σ*(*r*) matrix (blue) for the three Zinc-finger domain-containing proteins (A) REPIN1 (Q9BWE0), (B) Oxysterol receptor LXR-beta (P55055), and (C) Insulin gene enhancer (P61371). Simulation parameters were *t:* = 15 kJ mol*^−^*^1^, *α*_PAE_ = 15 nm*^−^*^1^, and *β*_PAE_ = 0.3 nm.

### Proteome-scale simulations with AF-CALVADOS

To demonstrate the scalability of AF-CALVADOS we performed AF-CALVADOS simulations of 12,483 full-length human proteins. Specifically, we selected all human proteins that (i) have lengths 30–1500 amino acid residues, and (ii) are predicted to be intracellular and without transmembrane regions (see Methods). We envisage that these conformational ensembles can be used to analyse sequence-dynamics-function relationships at scale, or can be used to train generative models in the same way as HPS simulations of IDRs have been used^27–29^.

To illustrate how folded domains may influence the conformational properties of IDRs, we focused on a set of 1,487 transcription factors in our dataset. For comparisons, we selected all IDR fragments with lengths of 30 or more amino-acid residues within each of these proteins (see Methods) and also performed simulations of these IDRs alone, for a total of 3,339 IDR segments. We compare a length-normalised average end-to-end distances 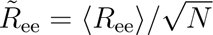 and internal-distance scaling exponents (*ν*) of the IDRs either in the context of the full-length transcription factor or as isolated IDR (Fig. 7A,D). We performed the analysis separately for IDR segments located at the N- or C-termini (‘terminal’) (Fig. 7B,E), and those located between folded domains (‘central’) (Fig. 7C,F). We find that *R̃*_ee_ and *ν* between the isolated IDRs and the IDR in a full-length context are generally correlated (Fig. 7A,D). For a subset of IDRs, in particular among ‘central’ IDRs located between folded domains, we find that the IDRs in context of the full-length proteins is substantially more compact than the isolated IDR (Fig. 7C,F); this substantial compaction in full-length context is more pronounced for shorter IDRs (SI Fig. S8). Short IDRs connecting folded regions are thus mostly affected by their MDP context whereas longer IDRs and terminal IDRs show more similar global dimensions on their own and in MDP context. We stress, however, that the spread around the diagonal corresponds to sizable changes in conformational properties with a magnitude of biological relevance^30,31^.

**Figure 7:**
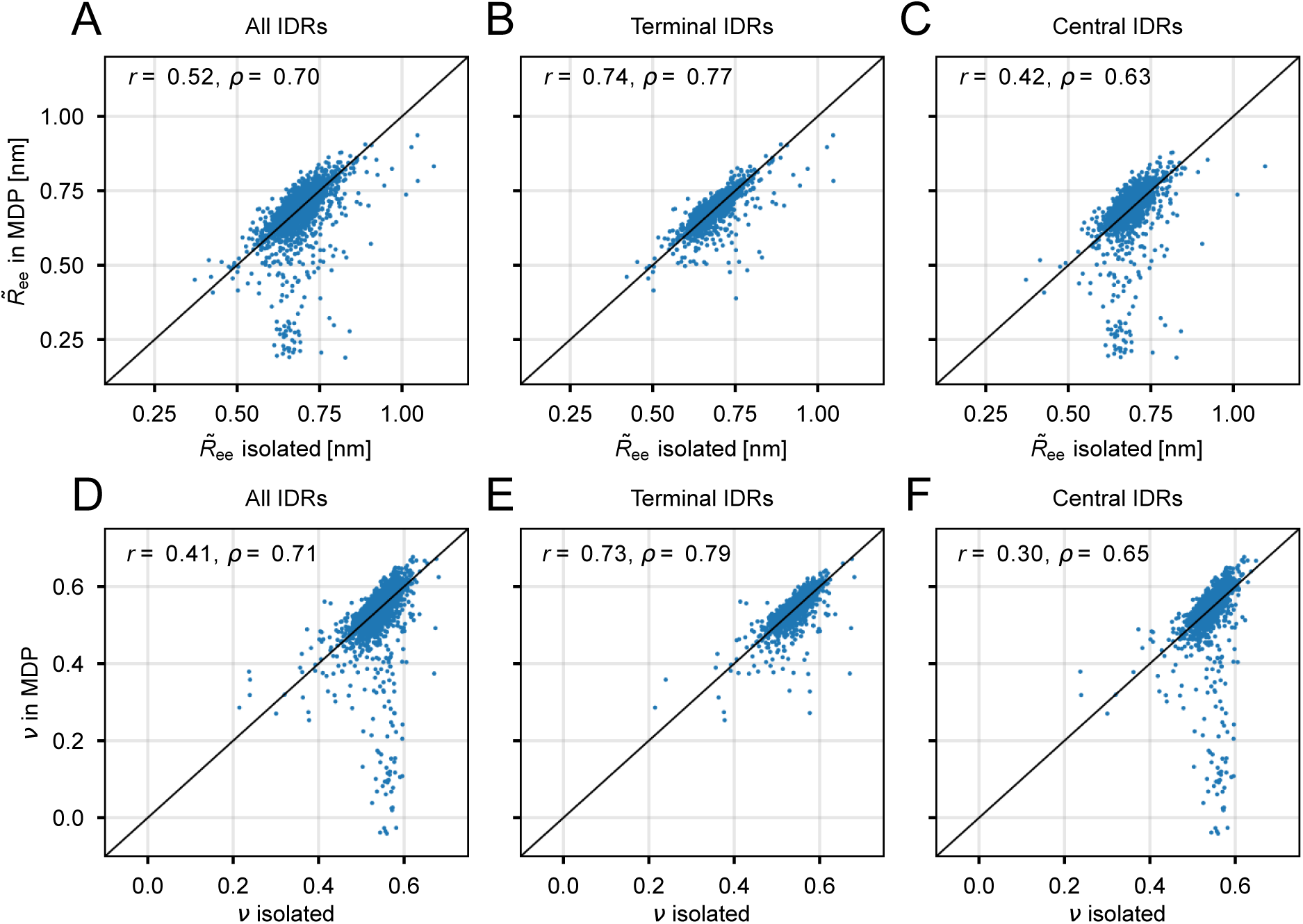
Comparison of IDRs in transcription factors either alone (‘isolated’) or in the context of the full-length protein (‘in MDP’). (A–C) Length-normalised average end-to-end distances 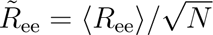 Internal-distance scaling exponents (*ν*). We show (A,D) the entire set of 3,339 IDRs in 1,487 transcription factors, (B,E) IDRs that are located at the N- or C-termini, and (C,F) IDRs located between folded domains.

We asked if the strong compaction for some ‘central’ IDRs is driven by interactions between the flanking folded domains. We therefore correlated the context-dependent differences Δ*R̃*_ee_ = *R̃*_ee_(in MDP) − *R̃*_ee_(isolated) and Δ*ν* = *ν*(in MDP) − *ν*(isolated) with the average simulation residue fluctuations *σ*(*r*) between the 20 residues preceding the IDR and the 20 residues following the IDR in sequence to probe whether those two groups of 20 residues move as a unit (SI Fig. S9). Most of the data are scattered around slight expansion in full-length context, with median Δ*R̃*_ee_ = 0.018 nm and median Δ*ν* = 0.009 (SI Fig. S9). However, some sequences show extreme compaction in the full-length context. For these IDRs, the flanking folded residues have strongly coupled motion (low average *σ*(*r*)) in the simulations. This suggests that strong IDR compaction in the full-length context is indeed driven by the correlated motion of the flanking folded residues. Visual inspection confirmed that extreme differences in the expansion of isolated IDRs and IDRs in context were due to the flanking residues belonging to the same domain, effectively rendering the IDRs as long loops (SI Fig. S10).

Finally, we correlated the context dependence of the IDR chain expansion with IDR sequence features other than sequence length (SI Figs. S11 and S12). We selected features that previously have been described to be useful to predict IDR single-chain properties and phase behaviour, and are described elsewhere^32^. In brief, *λ̅* describes the average chain stickiness, *f*_aro_ is the fraction of aromatic residues, SHD is a hydrophobicity patterning parameter (including the overall hydrophobicity) with higher SHD values indicating a more patchy distribution^33^, NCPR is the net charge per residue, FCR is the fraction of charged residues, SCD is a charge patterning parameter with lower (more negative) SCD values indicating patches of like charges^34^, AH_pairs_ is a hydrophobicity parameter considering amino-acid sizes^32^, and *ν*_SVR_ predicts the IDR single-chain expansion^8^. We note that the interpretation of the trends is complicated by the inherent correlation between the sequence features and with sequence length *N* and amino acid composition. For example, NCPR is correlated with FCR and SCD, and SHD is correlated with *N*, *f*_aro_ and *λ̅*, among others.

To focus on those IDRs whose compaction in full-length context is not explained by being a loop between two coupled folded (or the same folded) domains, we excluded IDR sequences from the analysis that showed strong coupled motion between their flanking regions (*σ*(*r*) *<* 1.0 nm) (SI Fig. S9). Overall, we observe subtle but consistent associations between the difference in length-normalised average end-to-end distance Δ*R̃*_ee_ and several sequence features, with the strongest trends visible for NCPR and SHD (SI Fig. S11). The same trends are visible for Δ*ν* (SI Fig. S12).

Negative NCPR values correlate with a somewhat stronger compaction of the IDR in the full-length context compared to the isolated chain, whereas positive NCPR values show a trend toward expansion in the full-length context (Figs. S11 and S12). We speculate that this behaviour may be due to attractive/repulsive interactions of the negatively/positively charged IDRs with positively charged DNA-binding domains prevalent in transcription factors, causing the negatively charged IDR chain to transiently fold back onto the folded domain in the MDP.

Low hydrophobic patchiness (SHD) of the IDR sequence is associated with chain expansion in the full-length context (Figs. S11 and S12). We assume that this trend is mostly from short (generally low SHD) sequences. Even though some short IDRs identified as loops show substantial compaction in the full-length context (see above), many of the remaining non-loop IDRs expand slightly in the full-length context (scatter above the identity lines in SI Fig. S8 for 30 ≤ *N <* 88). Further work will be required to more systematically quantify the subtle contributions of the correlated sequence features of the IDRs as well as the associated folded domains to the global chain properties in the full-length context.

### Conclusions

Most proteins contain mixtures of folded domains and disordered regions, and the interplay between structure and disorder is often key to biological function. These proteins fall between the possibilities afforded by machine-learning models for predictions of the structures of folded domains and computationally efficient simulation approaches for disordered regions. We developed AF-CALVADOS to bridge this gap in studies of these flexible MDPs, and with the idea of developing an approach that is both accurate and scalable. We combine the power of AF2 for predicting the structures of and interactions between folded domains with the CALVADOS model for studying flexible proteins. Validation of AF-CALVADOS using experimental SAXS data from the recently described PeptoneBench benchmark demonstrates its accuracy and robustness across a wide range of structural architectures.

Despite its robustness and accuracy, AF-CALVADOS has some limitations. First, because CALVADOS represents each amino acid by a single bead, it does not capture local structuring within disordered regions, and for this reason we did not apply the PeptoneDB-CS benchmark^21^. Second, while CALVADOS may capture some effects of varying the temperature, pH or ionic strength, these effects are not included in AF2. Third, some folded elements may only form in the context of interactions in the native cellular environment. For example, the N-terminal PNt domain of *Bordetella pertussis* pertactin is predicted to be highly structured by AF2, whereas SAXS and CD experiments show that it is disordered as a monomer in solution^35,36^; AF-CALVADOS, like bAIes^19^, will generally model such proteins as folded. Finally, we note that the set of AF-CALVADOS parameters chosen here is not necessarily optimal for all sets of MDPs. However, our bootstrapping analysis indicates that a wide range of parameters give similar results for the proteins studied here, indicating that we are not over-fitting to the proteins we analysed. Depending on their goals, we envisage that users may want to use the AF-CALVADOS architecture and adapt the restraint parameters depending on their needs, e.g., to restrain folded domains more robustly (larger *t:* and *β*_PAE_) or to overrule AF2 information when it disagrees with experimental evidence.

Despite these limitations, we envisage that AF-CALVADOS will enable a wide range of studies linking protein structure, dynamics and function. Our database of 12,483 full-length human proteins is freely available and may be used to study individual proteins, families of proteins (such as transcription factors) or to train generative models for protein ensembles.

## Methods

### The AF-CALVADOS force field

We used a modified version of the CALVADOS 3 force field, that we have termed AF-CALVADOS, to simulate MDPs. The CALVADOS family of force fields are coarse-grained hydrophobicity scale (HPS) models, in which one amino acid residue is represented with one simulation bead. The force field and software have been described elsewhere^37^, and we here describe key aspects related to the force field and how AF-CALVADOS differs from CALVADOS 3. Bonds between neighbouring residues are modelled with a harmonic potential,

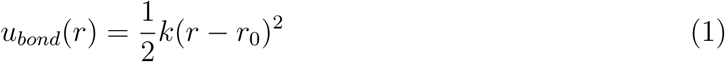

with *k* = 8033 kJmol*^−^*^1^nm*^−^*^2^ the force constant and *r*_0_ = 0.38 nm the equilibrium distance for unrestrained residues, with different values for restrained particle beads (see below).

Electrostatic interactions between nonbonded protein beads are modelled via a salt-screened Debye-Hückel (DH) term,

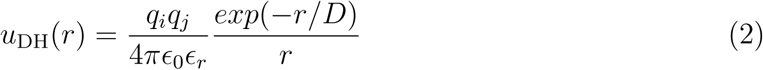

where *q_i_*is the charge of bead *i*, *t:*_0_ is the vacuum permittivity, 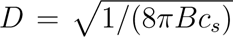 is the Debye length of an electrolyte solution of ionic strength *c_s_*, and *B*(*t:_r_*) is the Bjerrum length of temperature-dependent dielectric constant *t:_r_* ^38^,

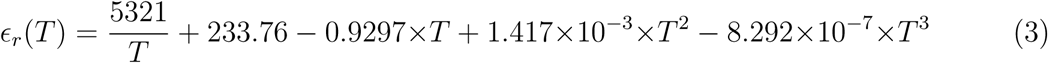

The potential is truncated and shifted with cutoff distance *r_c_* = 4 nm.

Non-electrostatic interactions between nonbonded protein beads are modelled via an Ashbaugh-Hatch (AH) potential^5^,

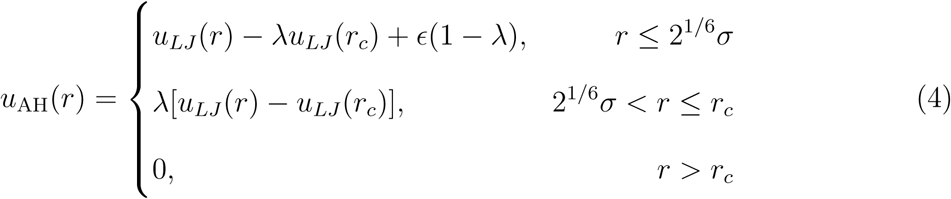

with Lennard-Jones (LJ) potential *u_LJ_* (*r*),

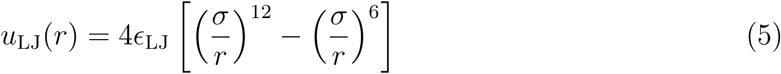

with *ε:*_LJ_ = 0.8368 kJ mol*^−^*^1^ and cutoff *r_c_* = 2.0 nm. The *λ* values are residue type-specific stickiness parameters optimised to reproduce single-chain properties of IDRs and MDPs well^10^.

Restraints in AF-CALVADOS are incorporated via a structure-based Gō-type potential. The restraining potential has the form:

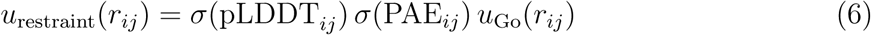

with the Gō potential defined as:

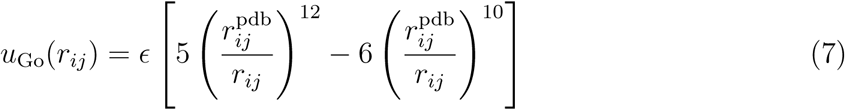

where *t:* is the tunable well depth of the Gō potential (in units of kJ/mol) and 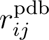 the centre-of-mass distance of residue pair *ij* in the starting structure. *σ*(pLDDT*_i,j_*) and *σ*(PAE*_i,j_*) encode AF2 information via the logistic functions,

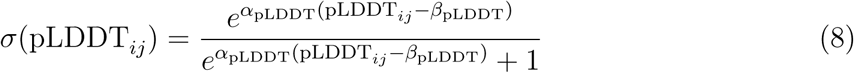

with pLDDT*_ij_*= min(pLDDT*_i_,* pLDDT*_j_*), and

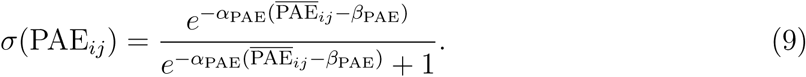

with 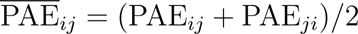 being the symmetrised PAE matrix values for residue pair *ij*.

The parameters *α*_pLDDT_ = 50 and *β*_pLDDT_ = 0.8 were fixed a priori to only restrain pairs for which the lower of the two pLDDT value is categorised as ‘high’ (0.7 *<* pLDDT*/*100 *<* 0.9) confidence, or ‘very high’ (pLDDT*/*100 *>* 0.9) confidence according to the AF2 classification^25^.

The parameters *t:*, *α*_PAE_, and *β*_PAE_ were tuned to reproduce experimental *R*_g_ data and to maintain the stability of folded domains as predicted by the PAE (see below). Only particle beads with distance *r_ij_ <*= 0.9 nm were restrained, in line with previous work^10^.

Contrary to CALVADOS 3, where domains are classified as folded or disordered in a binary manner, and as such are either restrained with uniform force constant or not restrained at all, the force constant for restraints varies in AF-CALVADOS. Therefore, edge cases including weak restraints need to be considered to avoid situations where AH and DH nonbonded interactions are turned off but the restraints are very weak, leading to artificially weak nonbonded interactions.

To remedy this, we gradually switch on AH and DH nonbonded (nb) interactions for weakly restrained residue pairs (*<*≈ 0.1 of full restraints) according to the logistic function:

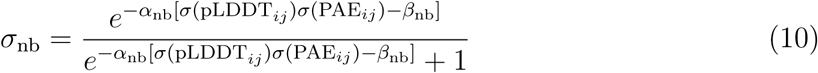

with *α*_nb_ = 80 and *β*_nb_ = 0.1.

For computational efficiency, for nonbonded pairs with very low restraining prefactor *σ*(pLDDT*_ij_*) *σ*(PAE*_ij_*) *<* 0.05, the restraint is turned off and the residues are treated like IDRs in CALVADOS 3 (with AH and DH potential). Bonded pairs with very low restraining prefactor are assigned an equilibrium bond distance *r*_0_ = 0.38 nm.

Conversely, for nonbonded pairs with a high restraining prefactor (*σ*(pLDDT*_ij_*) *σ*(PAE*_ij_*) *>* 0.15), only restraints are applied and AH and DH potentials are turned off. Bonded pairs with high restraining prefactor are assigned an equilibrium bond distance of the centre-of-mass distance in the starting structure 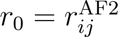.

For non-bonded pairs with intermediate restraining prefactor (0.05 *< σ*(pLDDT*_ij_*) *σ*(PAE*_ij_*) *<* 0.15), the potential between nonbonded restrained particles is a sum of the restraint potential *u*_restraint_ (eq. 6) and the scaled (eq. 10) original CALVADOS 3 AH and DH potentials:

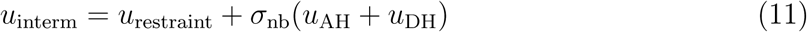

Bonded pairs with intermediate restraining prefactor have equilibrium bond length

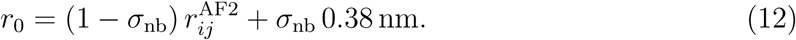

### Intracellular dataset

We selected human intracellular protein sequences from the human proteome of the AF2 database (AFDB) version 4^39^. We extracted sequences from the SEQRES header section of PDB files. For 19,447 non-redundant sequences of length 30 to 1,500, we predicted transmembrane (TM) regions using DeepTMHMM^40^ via biolib and selected 12,483 sequences predicted as intracellular/cytosolic (all residues state ‘I’).

Among the intracellular proteins we annotated transcription factors based on a published list of transcription factor genes^41^ and a map of these to Uniprot entries^42^.

### Molecular dynamics simulations

For the *R*_g_-based parameterisation of AF-CALVADOS, we performed simulations of the 23 MDP proteins from Cao et al.^10^ in a parameter grid scan with *t:* = {5, 10, 15, 20, 25} kJ mol*^−^*^1^, *α*_PAE_ = {15, 30} nm*^−^*^1^, and *β*_PAE_ = {0.1, 0.2, 0.3, 0.4, 0.5} nm.

We simulated each protein with a Langevin integrator with integration time step *t* = 10 fs and friction coefficient *ν* = 0.01 ps*^−^*^1^ in a constant volume (NVT) ensemble in a cubic simulation box with edge size 80 nm and periodic boundary conditions at experimental temperature, ionic strength, and pH value, as listed in Tables S3 and S5 in Cao et al. ^10^.

We sampled 2010 conformations per protein and parameter set, and discarded the first 10 conformations as equilibration. In order to sample uncorrelated structures, we recorded the conformations at intervals *w* scaled by the sum of ‘disordered’ (fully unrestrained) residues

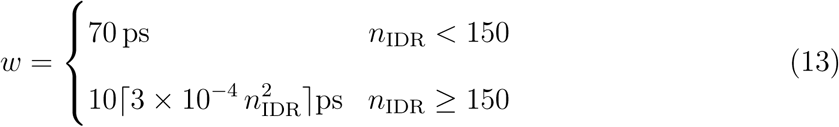

where […] denotes the ceiling function^9^.

To examine whether folded domains are stable in simulations at 310 K, we randomly drew a set of 30 entries from the dataset of intracellular proteins described above, downloaded AF2 structure and PAE files from the EBI-AF database, and performed simulations at different parameters *t:*, *α*_PAE_, and *β*_PAE_ as before, except for a reduction to 1010 sampled conformations and fixed conditions *T* = 310 K, *I* = 0.15 M, and *pH* = 7.0. For these proteins, we calculated the Pearson correlation coefficient *r* and Spearman correlation coefficient *ρ* between the flattened PAE and *σ*(*r*) matrices for all parameter combinations. To equalise the contributions from low and high values in each PAE matrix to the correlation coefficient (most of the values for IDPs are high, and most of the values for predominantly ordered proteins are low), we balanced the datasets per protein by binning the data into 10 equidistant bins from lowest to highest PAE value, repeatedly drawing 1000 samples each from each bin and re-analysing the correlation coefficients for the resampled, balanced data. The reported correlation coefficients *r* and *ρ* are averages for 100 repeats of the above procedure. Example correlation plots of the unbalanced datasets for two parameter sets are shown in SI Fig. S13.

The 12,483 proteome entries were extracted from AFDB and simulated at *T* = 310 K, *I* = 0.15 M, and *pH* = 7.4 to generate 1010 frames. For all transcription factors, we also extracted all IDRs based on the criterion of contiguously fully unrestrained stretches of sequences of length 30 residues or greater. The individual IDRs in the transcription factors were simulated with the same simulation settings as the full-length proteins.

### Benchmarking using PeptoneDB-SAXS

For each of the 439 sequences in the PeptoneDB-SAXS benchmark^21^, we generated 1000 conformations using AF-CALVADOS at the indicated pH and salt concentration, and using previously generated AF2 structures, pLDDT and PAE scores^21^. We converted the coarse grained AF-CALVADOS structures to all-atom representations using cg2all^43^ (with settings –cg ResidueBasedModel). We calculated SAXS curves using Pepsi-SAXS^44^ with parameters from and wrapper scripts released with PeptoneBench^21^. Pepsi-SAXS did not produce output for any structures of SASDRS5, and we further removed eight proteins for which the SAXS data were measured in 1–6 M urea because the effect of urea is not considered in any of the models. Of the remaining 430 proteins, 20 had ≥ 80% sequence identity (global alignment) to the proteins used in the development of CALVADOS 3 or AF-CALVADOS and were marked as training data to ensure a fairer comparison. The root-mean-square-error and reweighting calculations were carried out as in the original PeptoneBench implementation with a target effective sample size of 10 conformations. For the most direct comparison to the other methods (Fig. 5) we use 100 conformations (sampled as every tenth of the 1000 structures); the accuracy of AF-CALVADOS is slightly greater when we use all 1000 conformations (SI Fig. S4). In the benchmark, the label AlphaFold represents the single conformation used as input to AF-CALVADOS and is only shown for reference. PeptoneBench includes a filter for unphysical conformations at the all-atom level, mostly relevant for comparison to the calculation of NMR chemical shifts. Application of this filter reduces the ensemble size substantially, and we provide comparisons of the results with (SI Fig. S14) or without this filter (Fig. 5). The disorder G-scores from ADOPT2 and conformational ensembles from other methods were downloaded from the PeptoneBench material (https://github.com/PeptoneLtd/PeptoneBench). Error bars for each LOWESS summary curve are the 5 and 95 percentile of 200 bootstrap LOWESS regressions with the same number of proteins.

## Data and Code Availability

Data and code used for this work is available as a GitHub repository: https://github.com/KULL-Centre/_2025_buelow_AF-CALVADOS. PeptoneBench is available from https://github.com/PeptoneLtd/PeptoneBench. Protein structure and simulation data are available as an Electronic Research Data Archive (ERDA) at the University of Copenhagen repository: https://sid.erda.dk/sharelink/gfZNiWkxRT.

## Acknowledgments

We thank Michele Invernizzi, Sandro Bottaro, Kamil Tamiola and other members of the Peptone Ltd. team for discussions and help regarding PeptoneBench. We acknowledge access to computational resources from the ROBUST Resource for Biomolecular Simulations (supported by the Novo Nordisk Foundation grant no. NF18OC0032608). This work is a contribution from the PRISM (Protein Interactions and Stability in Medicine and Genomics) centre funded by the Novo Nordisk Foundation (to K.L.-L.; NNF18OC0033950).

## Supplementary Information

**Figure S1:**
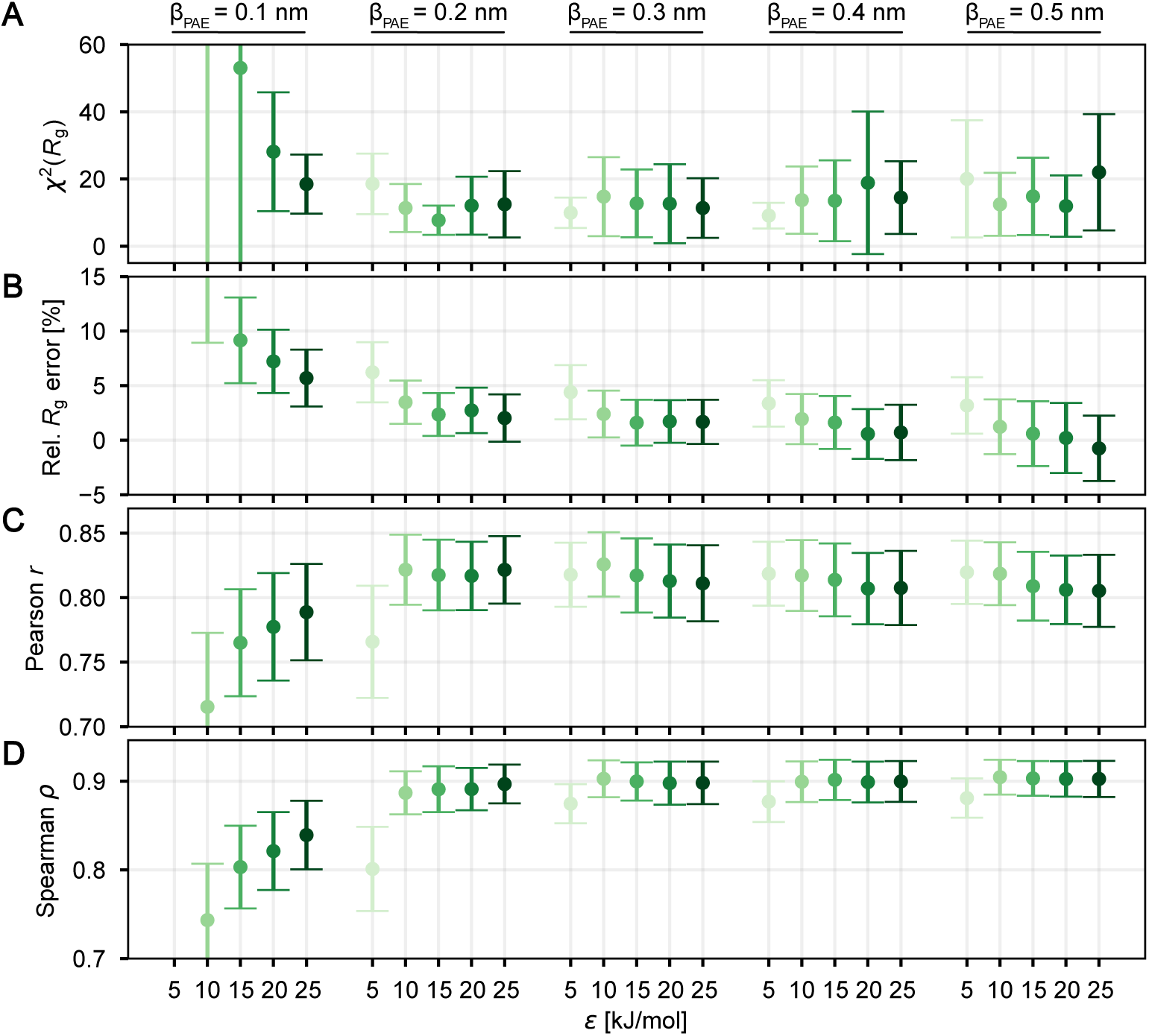
AF-CALVADOS parameterisation with *α*_PAE_ = 30 nm*^−^*^1^ vs. experimental *R*_g_ data (A, B) and using metrics describing the recovery of AF2 PAE matrix (C, D). Error bars indicate 95% confidence intervals estimated by 1000-fold bootstrapping.

**Figure S2:**
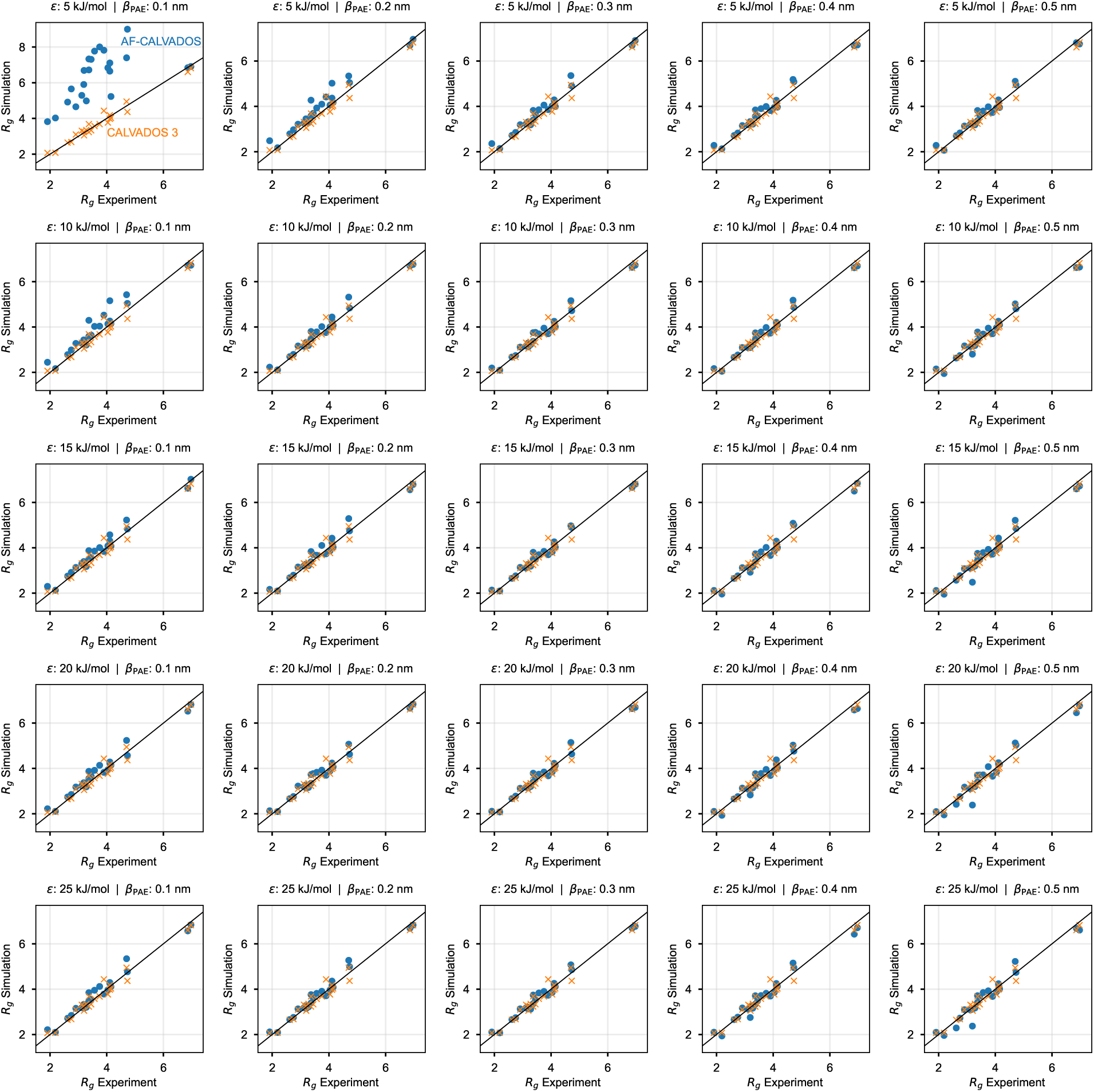
Comparison of *R*_g_ estimated from simulation vs. experimental *R*_g_ for restraining parameter scan (*α*_PAE_ = 15 nm*^−^*^1^).

**Figure S3:**
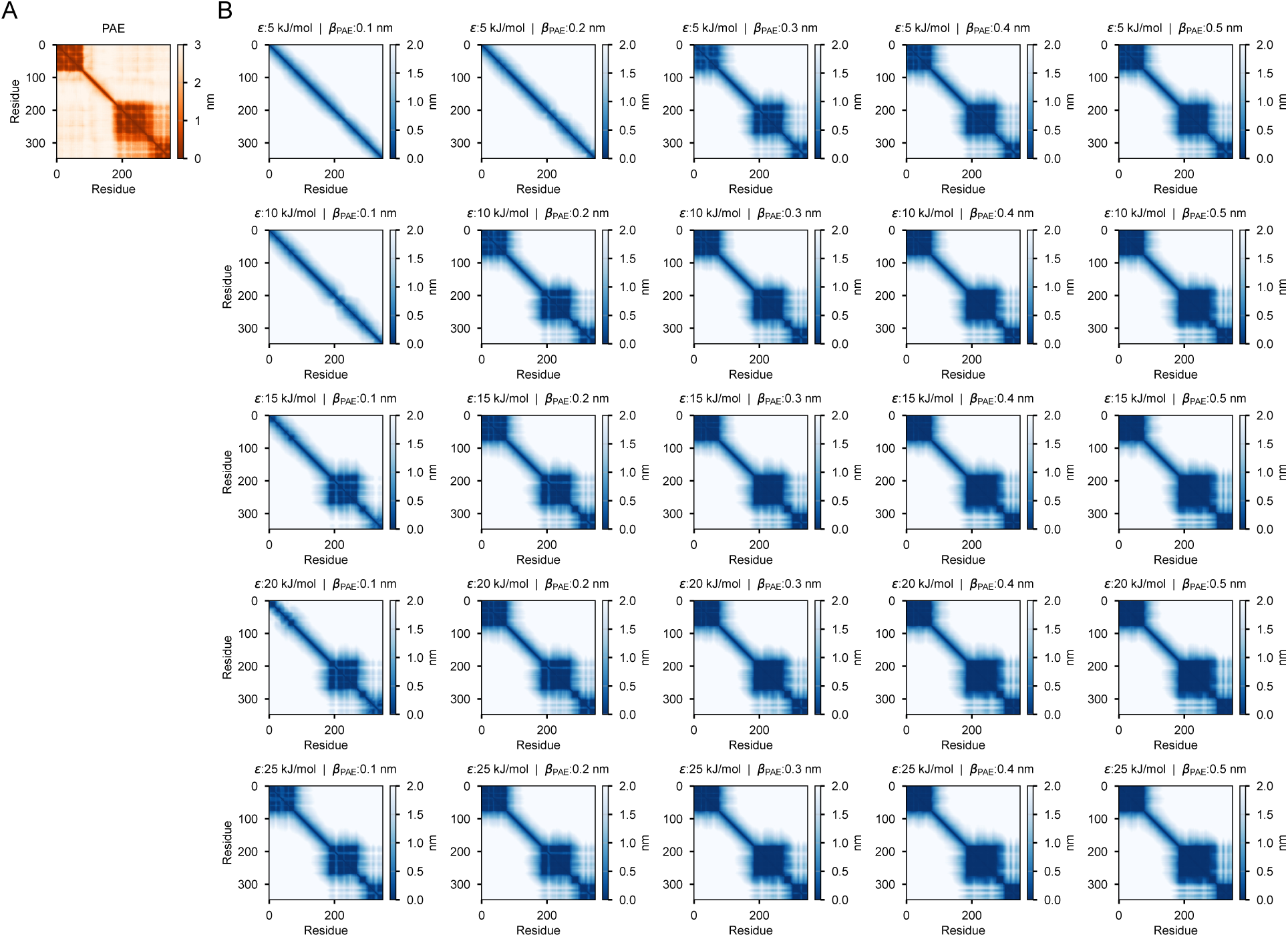
Comparison of the PAE matrix (A) vs. set of *σ*(*r*) matrices (B) for transcription elongation factor TCEA3 (Uniprot entry: O75764).

**Figure S4:**
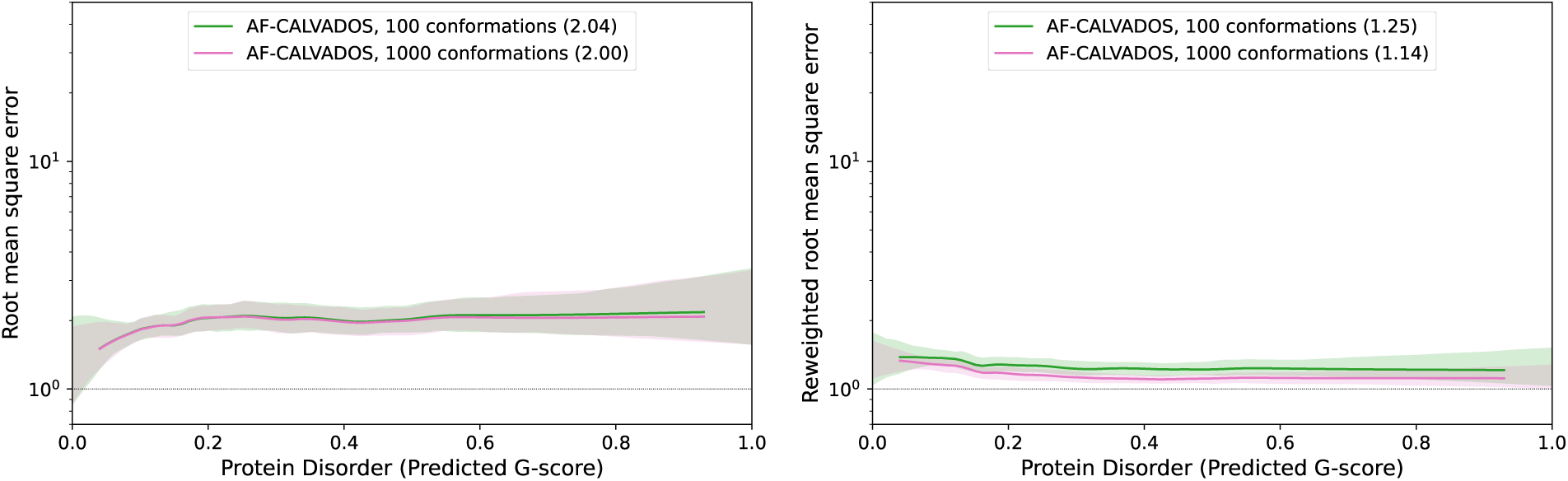
Comparison of summary statistics of the PeptoneDB-SAXS benchmark for AF-CALVADOS ensembles with 100 and 1000 structures. (Left) The larger ensemble improves the performance slightly. (Right) When the ensembles are scored after reweighting to the data, the difference becomes larger as the reweighting can ‘select’ among a more diverse set of conformations.

**Figure S5:**
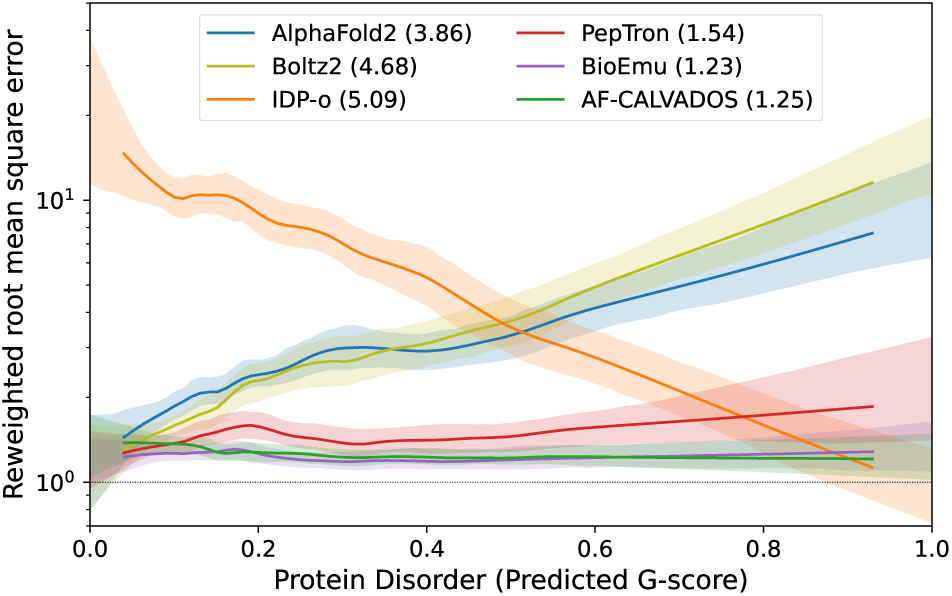
Summary statistics of the PeptoneDB-SAXS benchmark when combined with reweighting. The area under the curve is given in parenthesis after each method.

**Figure S6:**
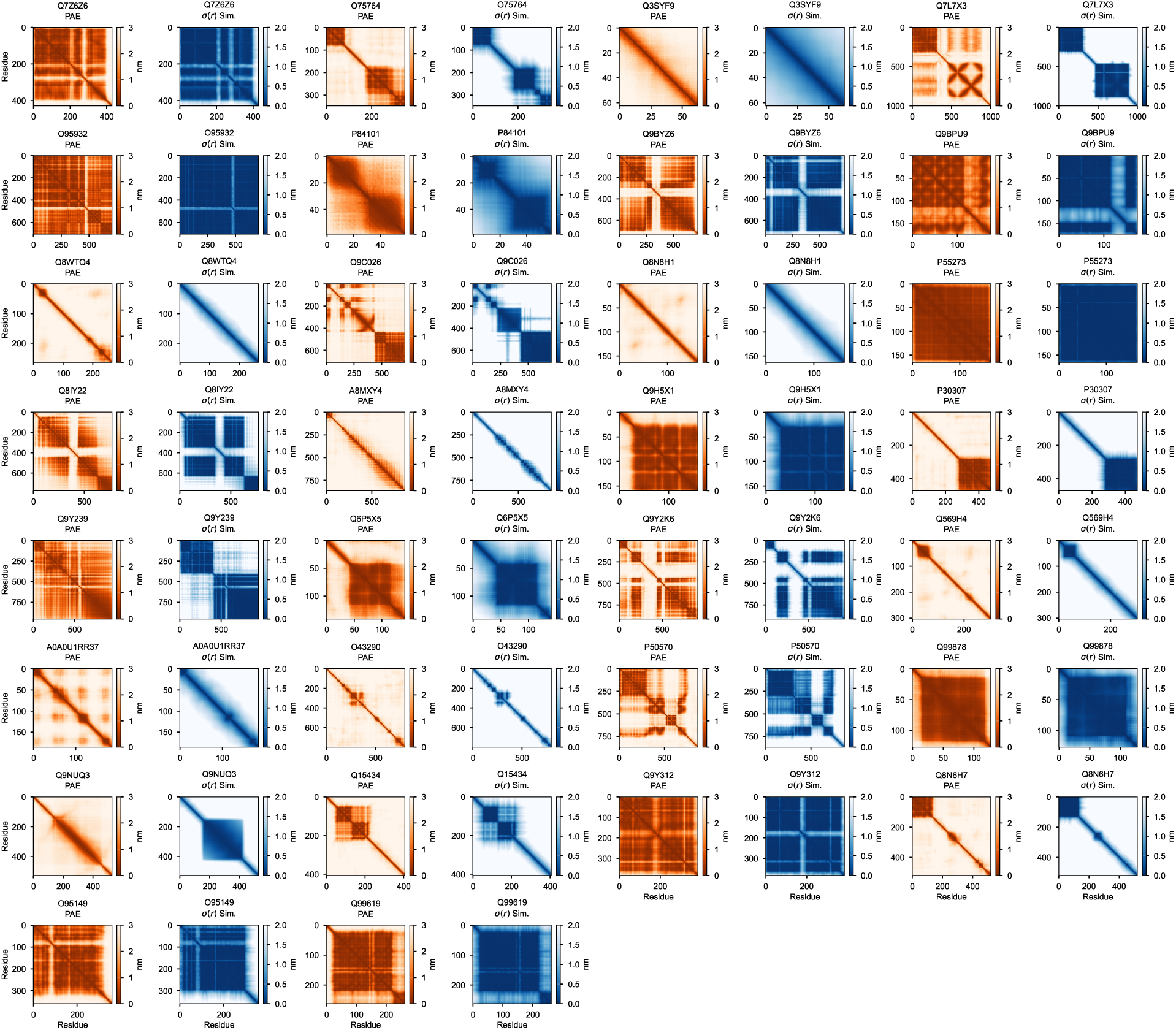
Comparison of input AF2 PAE matrix with standard deviation of residue pair distances *σ*(*r*) for the 30 intracellular proteins used for parameterisation.

**Figure S7:**
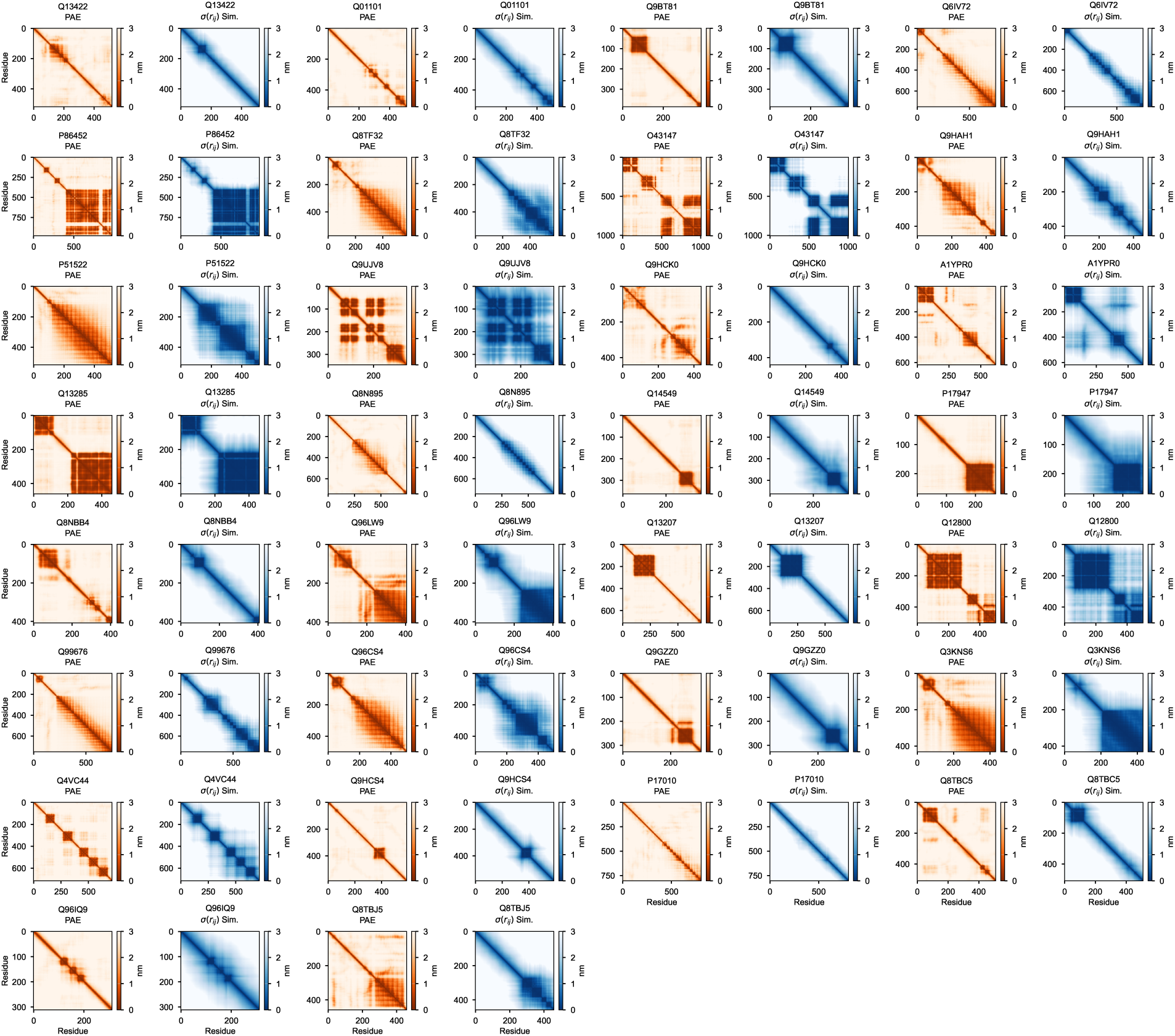
Comparison of input AF2 PAE matrix with standard deviation of residue pair distances *σ*(*r*) for 30 randomly selected transcription factors from the set of intracellular proteins that were not used in parameterisation.

**Figure S8:**
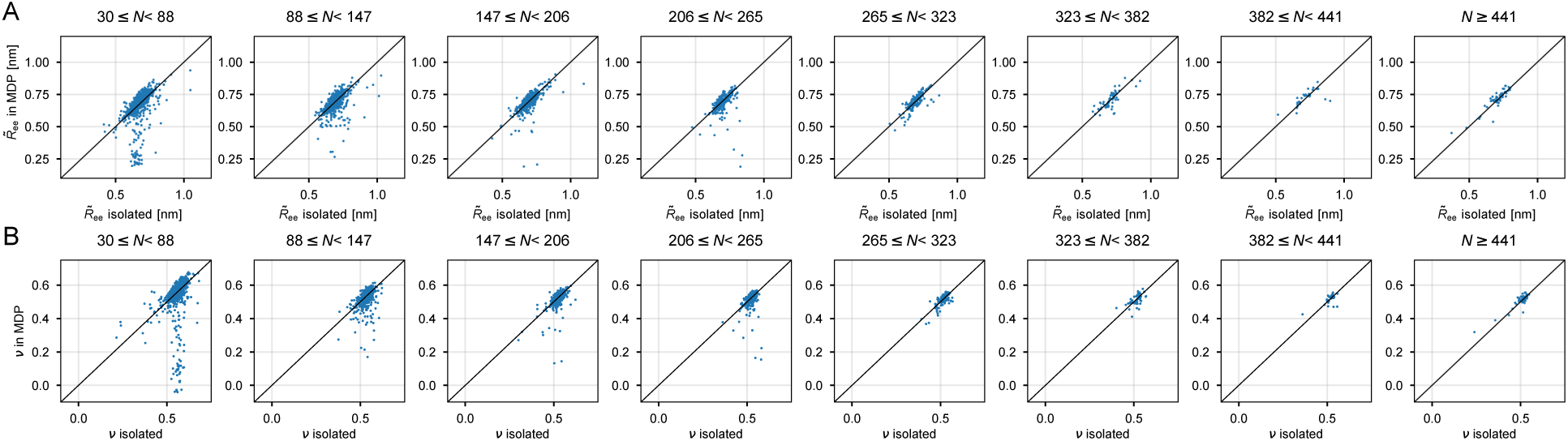
Comparison of disordered regions from transcription factors either as isolated IDR or IDRs in their full-length MDP context. The data are broken down for different chain lengths (*N*). (A) Length-normalised average end-to-end distances and (B) internal-distance scaling exponents.

**Figure S9:**
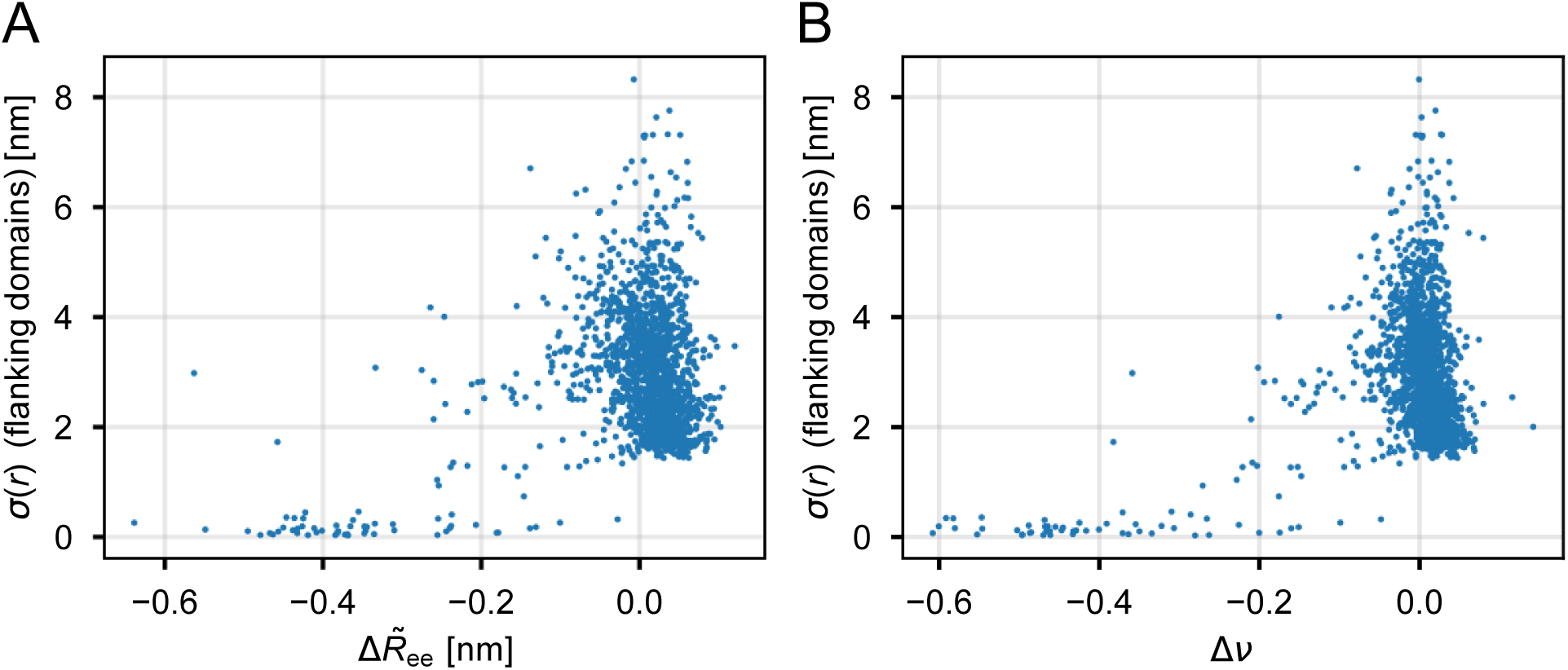
Differences in transcription factor expansion in MDP context and isolation vs. the average residue distance fluctuations *σ*(*r*) between the 20 flanking residues of the IDR (i.e. between the 20 residues before and 20 residues after the IDR). (A) Length-normalised average end-to-end distance difference Δ*R̃*_ee_ = *R̃*_ee_(in MDP) − *R̃*_ee_(isolated). (B) Internal-distance scaling exponent difference Δ*ν* = *ν*(in MDP) − *ν*(isolated).

**Figure S10:**
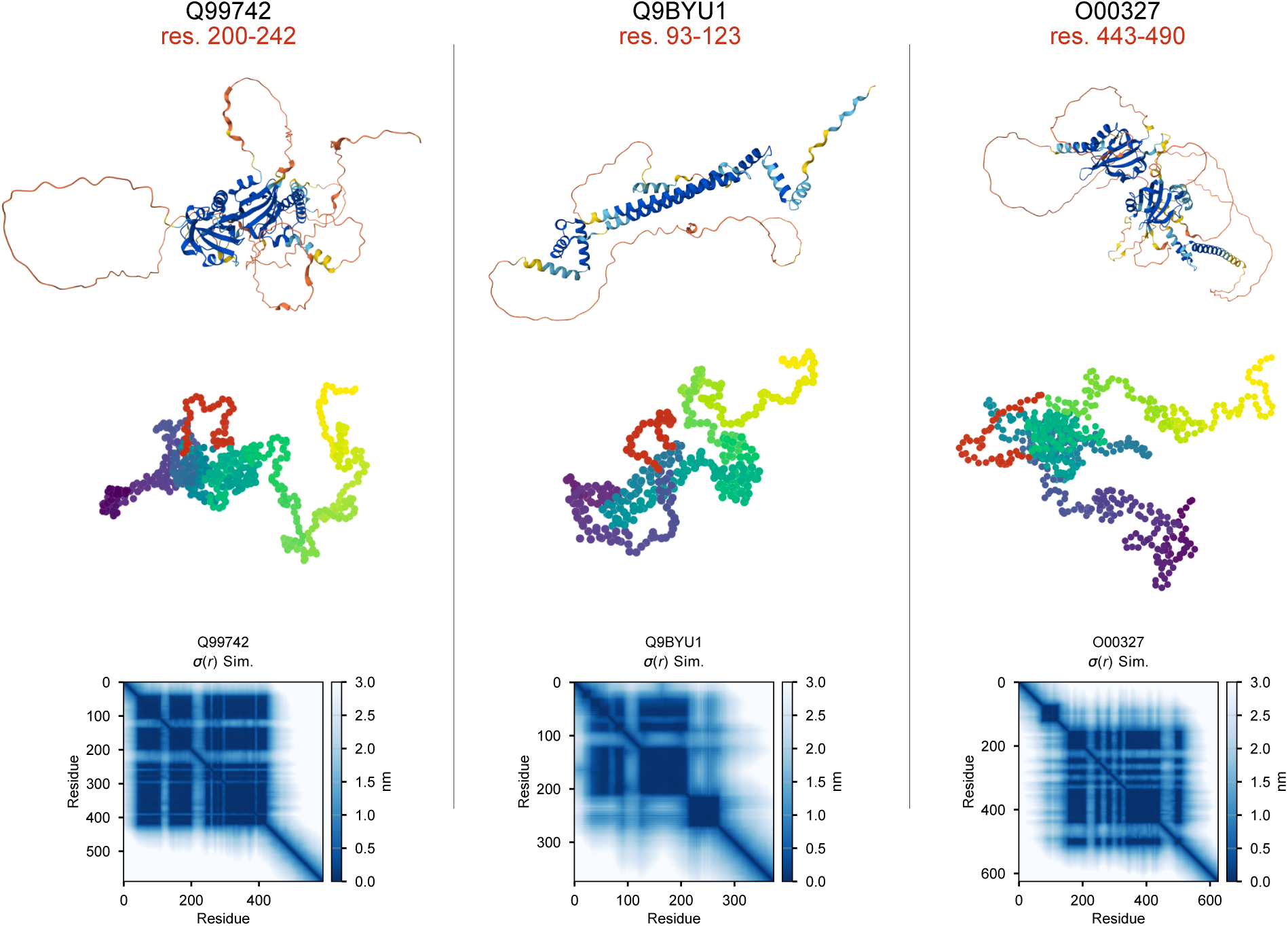
Examples of loop structures corresponding to large negative deviation of IDR chain expansion in MDP context. The loop region is coloured in red. Top row: EBI AlphaFold2 structure prediction. Middle row: Simulation snapshot. Bottom row: *σ*(*r*) matrix.

**Figure S11:**
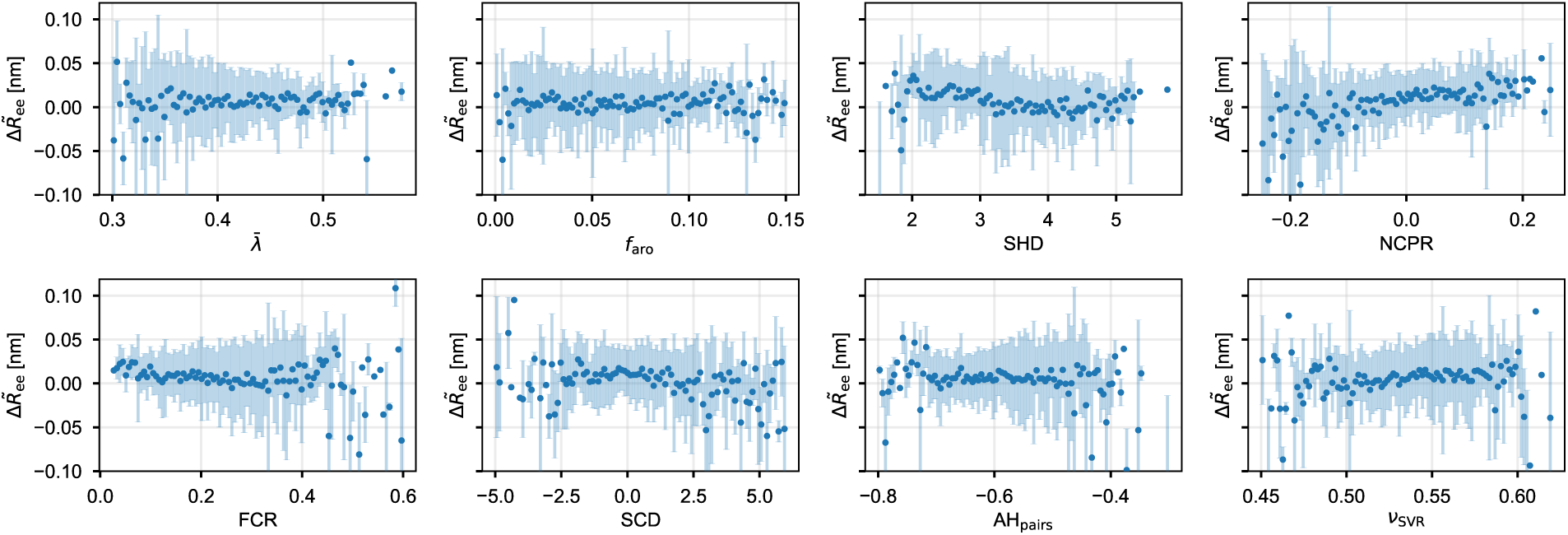
Binned length-normalised average end-to-end distance difference Δ*R̃*_ee_ = *R̃*_ee_(in MDP) *R̃*_ee_(isolated) vs. IDR sequence features. IDR sequences with strongly interacting flanking regions (*σ*(*r*) *<* 1.0 nm) are excluded. Error bars show standard deviations in each bin.

**Figure S12:**
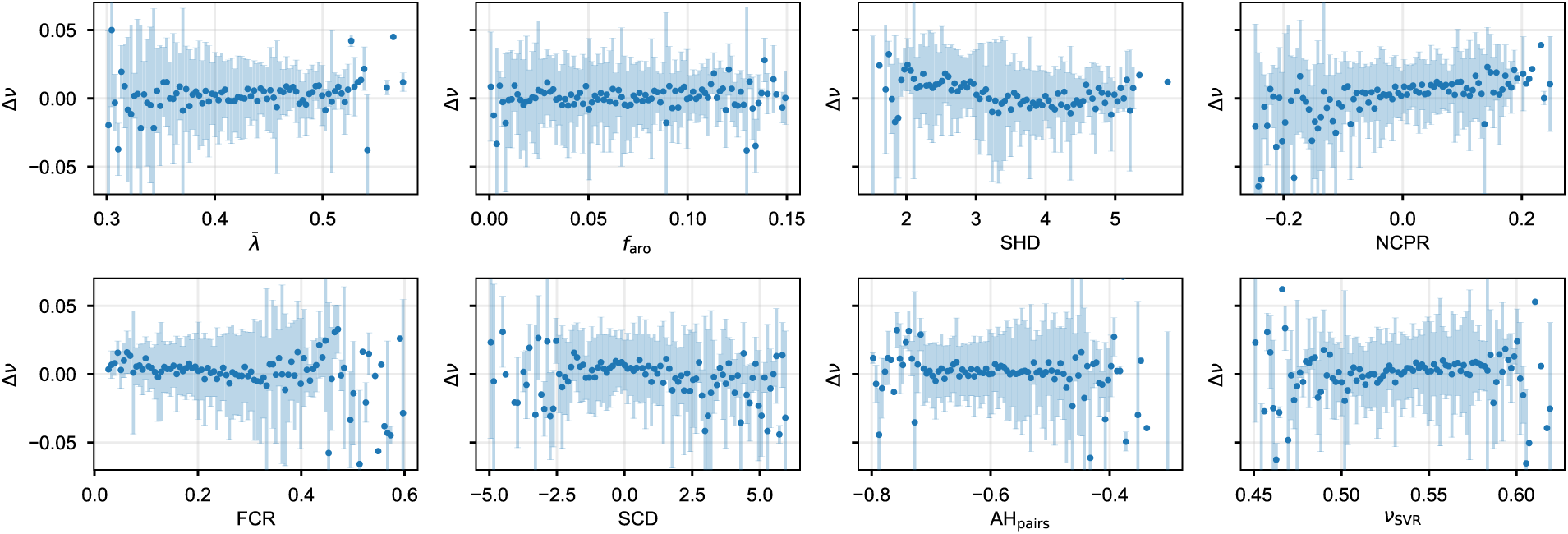
Binned internal-distance scaling exponent difference Δ*ν* = *ν*(in MDP) *ν*(isolated) vs. IDR sequence features. IDR sequences with strongly interacting flanking regions (*σ*(*r*) *<* 1.0 nm) are excluded. Error bars show standard deviations in each bin.

**Figure S13:**
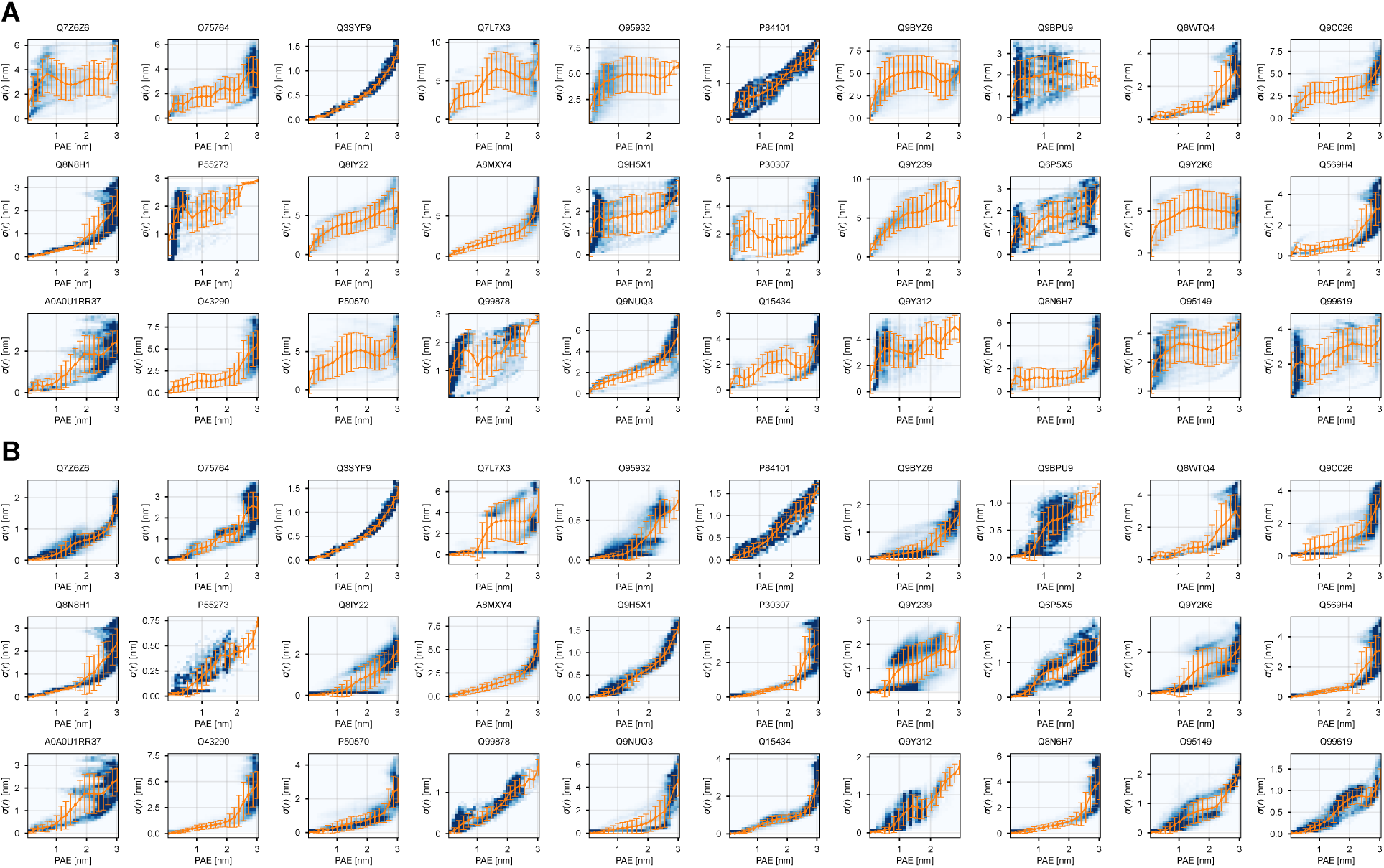
Correlation between flattened PAE and *σ*(*r*) values. Blue shading indicates point density in a 2D histogram with 30 30 bins. Orange plots show mean and standard deviation of values binned across PAE values (*n* = 20 bins). All plots are shown with *α*_PAE_ = 15 nm*^−^*^1^. (A) *t:* = 5 kJ mol*^−^*^1^ and *β*_PAE_ = 0.1 nm. (B) *t:* = 15 kJ mol*^−^*^1^ and *β*_PAE_ = 0.3 nm.

**Figure S14:**
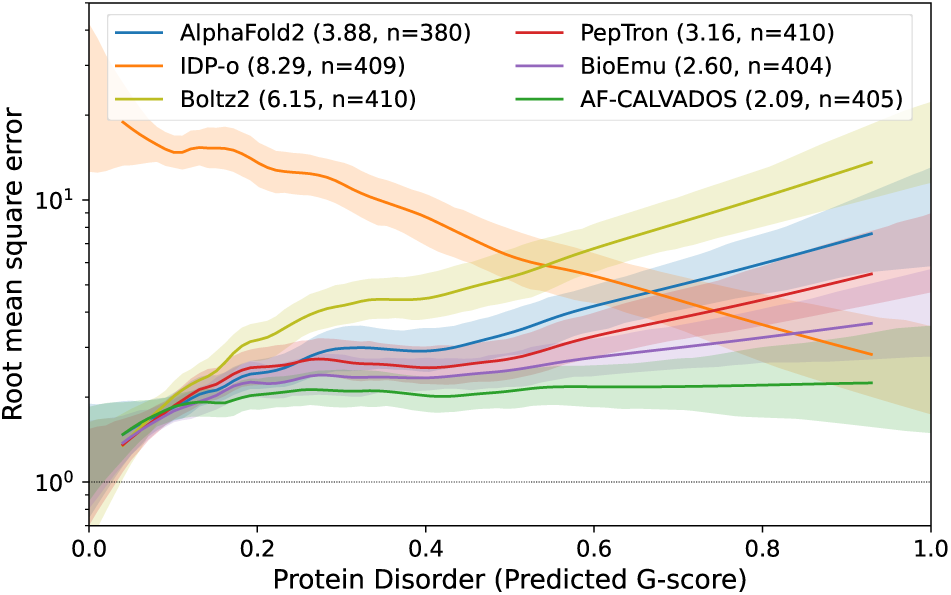
Summary statistics of the PeptoneDB-SAXS benchmark where the filter of unphysical conformations from PeptoneBench has been applied. The area under the curve is given in parenthesis after each method followed by the number of proteins for which all conformations have not been filtered out.

## References

(1) Tunyasuvunakool, K. et al. Highly Accurate Protein Structure Prediction for the Human Proteome. Nature 2021, 596, 590–596.

(2) von Bülow, S.; Tesei, G.; Lindorff-Larsen, K. Machine Learning Methods to Study Sequence–Ensemble–Function Relationships in Disordered Proteins. Current Opinion in Structural Biology 2025, 92, 103028.

(3) Han, J.-H.; Batey, S.; Nickson, A. A.; Teichmann, S. A.; Clarke, J. The Folding and Evolution of Multidomain Proteins. Nat Rev Mol Cell Biol 2007, 8, 319–330.

(4) Tsang, B.; Pritišanac, I.; Scherer, S. W.; Moses, A. M.; Forman-Kay, J. D. Phase Separation as a Missing Mechanism for Interpretation of Disease Mutations. Cell 2020, 183, 1742–1756.

(5) Ashbaugh, H. S.; Hatch, H. W. Natively Unfolded Protein Stability as a Coil-to-Globule Transition in Charge/Hydropathy Space. J. Am. Chem. Soc. 2008, 130, 9536–9542.

(6) Dignon, G. L.; Zheng, W.; Kim, Y. C.; Best, R. B.; Mittal, J. Sequence Determinants of Protein Phase Behavior from a Coarse-Grained Model. PLOS Computational Biology 2018, 14, e1005941.

(7) Tesei, G.; Schulze, T. K.; Crehuet, R.; Lindorff-Larsen, K. Accurate Model of Liquid–Liquid Phase Behavior of Intrinsically Disordered Proteins from Optimization of Single-Chain Properties. Proceedings of the National Academy of Sciences 2021, 118, e2111696118.

(8) Tesei, G.; Trolle, A. I.; Jonsson, N.; Betz, J.; Knudsen, F. E.; Pesce, F.; Johansson, K. E.; Lindorff-Larsen, K. Conformational Ensembles of the Human Intrinsically Disordered Proteome. Nature 2024, 626, 897–904.

(9) Tesei, G.; Lindorff-Larsen, K. Improved Predictions of Phase Behaviour of Intrinsically Disordered Proteins by Tuning the Interaction Range [Version 2; Peer Review: 2 Approved]. Open Research Europe 2023, 2.

(10) Cao, F.; von Bülow, S.; Tesei, G.; Lindorff-Larsen, K. A Coarse-Grained Model for Disordered and Multi-Domain Proteins. Protein Science 2024, 33, e5172.

(11) McBride, A. C.; Yu, F.; Cheng, E. H.; Mpouli, A.; Soe, A. C.; Hammel, M.; Montelione, G. T.; Oas, T. G.; Tsutakawa, S. E.; Donald, B. R. Predicting pose distribution of protein domains connected by flexible linkers is an unsolved problem. Proteins: Structure, Function, and Bioinformatics 2025, 1–19.

(12) Akdel, M. et al. A Structural Biology Community Assessment of AlphaFold2 Applications. Nat Struct Mol Biol 2022, 29, 1056–1067.

(13) Guo, H.-B.; Perminov, A.; Bekele, S.; Kedziora, G.; Farajollahi, S.; Varaljay, V.; Hinkle, K.; Molinero, V.; Meister, K.; Hung, C.; Dennis, P.; Kelley-Loughnane, N.; Berry, R. AlphaFold2 Models Indicate That Protein Sequence Determines Both Structure and Dynamics. Sci Rep 2022, 12, 10696.

(14) Jussupow, A.; Kaila, V. R. I. Effective Molecular Dynamics from Neural Network-Based Structure Prediction Models. J. Chem. Theory Comput. 2023, 19, 1965–1975.

(15) Brotzakis, Z. F.; Zhang, S.; Murtada, M. H.; Vendruscolo, M. AlphaFold Prediction of Structural Ensembles of Disordered Proteins. Nat Commun 2025, 16, 1632.

(16) Fernandes, N. P.; Gomes, T.; Cordeiro, T. N. Ensemblify: A User-Friendly Tool for Generating Ensembles of Intrinsically Disordered Regions of AlphaFold and User-Defined Models. bioRxiv 2025, 10.1101/2025.08.26.672300.

(17) Pajkos, M.; Clerc, I.; Zanon, C.; Bernadó, P.; Cortés, J. AFflecto: A web server to generate conformational ensembles of flexible proteins from AlphaFold models. Journal of Molecular Biology 2025, 169003.

(18) Wróblewski, K.; Kmiecik, S. Integrating AlphaFold pLDDT Scores into CABS-flex for enhanced protein flexibility simulations. Computational and Structural Biotechnology Journal 2024, 23, 4350–4356.

(19) Schnapka, V.; Morozova, T.; Sen, S.; Bonomi, M. Atomic Resolution Ensembles of Intrinsically Disordered and Multi-Domain Proteins with Alphafold. bioRxiv 2025, 10.1101/2025.08.26.672300.

(20) Lewis, S. et al. Scalable Emulation of Protein Equilibrium Ensembles with Generative Deep Learning. Science 2025, 389, eadv9817.

(21) Invernizzi, M.; Bottaro, S.; Streit, J. O.; Trentini, B.; Venanzi, N. A. E.; Reidenbach, D.; Lee, Y.; Dallago, C.; Sirelkhatim, H.; Jing, B.; Airoldi, F.; Lindorff-Larsen, K.; Fisicaro, C.; Tamiola, K. Advancing Protein Ensemble Predictions across the Order–Disorder Continuum. bioRxiv 2025, 10.1101/2025.10.18.680935.

(22) Zhang, O.; Liu, Z. H.; Forman-Kay, J. D.; Head-Gordon, T. Deep Learning of Proteins with Local and Global Regions of Disorder. arXiv 2025, 10.48550/arXiv.2502.11326.

(23) Liu, Z. H.; Zhang, O.; De Castro, S.; Sun, K.; Ghafouri, H.; Attafi, O. A.; Fawzi, N. L.; Tosatto, S. C.; Monzon, A. M.; Moses, A. M., et al. AlphaFlex: Ensembles of the human proteome representing disordered regions. bioRxiv 2025, 10.1101/2025.11.24.690279.

(24) Kikhney, A. G.; Borges, C. R.; Molodenskiy, D. S.; Jeffries, C. M.; Svergun, D. I. SASBDB: Towards an automatically curated and validated repository for biological scattering data. Protein Science 2020, 29, 66–75.

(25) Jumper, J. et al. Highly Accurate Protein Structure Prediction with AlphaFold. Nature 2021, 596, 583–589.

(26) Passaro, S.; Corso, G.; Wohlwend, J.; Reveiz, M.; Thaler, S.; Somnath, V. R.; Getz, N.; Portnoi, T.; Roy, J.; Stark, H., et al. Boltz-2: Towards accurate and efficient binding affinity prediction. bioRxiv 2025, 10.1101/2025.06.14.659707.

(27) Janson, G.; Valdes-Garcia, G.; Heo, L.; Feig, M. Direct Generation of Protein Conformational Ensembles via Machine Learning. Nat Commun 2023, 14, 774.

(28) Zhu, J.; Li, Z.; Zheng, Z.; Zhang, B.; Zhong, B.; Bai, J.; Hong, X.; Wang, T.; Wei, T.; Yang, J.; Chen, H.-F. Accurate Generation of Conformational Ensembles for Intrinsically Disordered Proteins with IDPFold. Advanced Science 2025, e11636.

(29) Novak, B.; Lotthammer, J. M.; Emenecker, R. J.; Holehouse, A. S. Accurate Predictions of Conformational Ensembles of Disordered Proteins with STARLING. bioRxiv 2025, 10.1101/2025.02.14.638373.

(30) Martin, E. W.; Holehouse, A. S.; Peran, I.; Farag, M.; Incicco, J. J.; Bremer, A.; Grace, C. R.; Soranno, A.; Pappu, R. V.; Mittag, T. Valence and Patterning of Aromatic Residues Determine the Phase Behavior of Prion-like Domains. Science 2020, 367, 694–699.

(31) González-Foutel, N. S. et al. Conformational Buffering Underlies Functional Selection in Intrinsically Disordered Protein Regions. Nat Struct Mol Biol 2022, 1–10.

(32) von Bülow, S.; Tesei, G.; Zaidi, F. K.; Mittag, T.; Lindorff-Larsen, K. Prediction of Phase-Separation Propensities of Disordered Proteins from Sequence. Proceedings of the National Academy of Sciences 2025, 122, e2417920122.

(33) Zheng, W.; Dignon, G.; Brown, M.; Kim, Y. C.; Mittal, J. Hydropathy Patterning Complements Charge Patterning to Describe Conformational Preferences of Disordered Proteins. J. Phys. Chem. Lett. 2020, 11, 3408–3415.

(34) Sawle, L.; Ghosh, K. A Theoretical Method to Compute Sequence Dependent Configurational Properties in Charged Polymers and Proteins. J. Chem. Phys. 2015, 143, 085101.

(35) Riback, J. A.; Bowman, M. A.; Zmyslowski, A. M.; Knoverek, C. R.; Jumper, J. M.; Hinshaw, J. R.; Kaye, E. B.; Freed, K. F.; Clark, P. L.; Sosnick, T. R. Innovative Scattering Analysis Shows That Hydrophobic Disordered Proteins Are Expanded in Water. Science 2017, 358, 238–241.

(36) Bowman, M. A.; Riback, J. A.; Rodriguez, A.; Guo, H.; Li, J.; Sosnick, T. R.; Clark, P. L. Properties of Protein Unfolded States Suggest Broad Selection for Expanded Conformational Ensembles. Proceedings of the National Academy of Sciences 2020, 117, 23356–23364.

(37) von Bülow, S.; Yasuda, I.; Cao, F.; Schulze, T. K.; Trolle, A. I.; Rauh, A. S.; Crehuet, R.; Lindorff-Larsen, K.; Tesei, G. Software Package for Simulations Using the Coarse-Grained CALVADOS Model. arXiv 2025, 10.48550/arXiv.2504.10408.

(38) Akerlof, G. C.; Oshry, H. I. The Dielectric Constant of Water at High Temperatures and in Equilibrium with Its Vapor. J. Am. Chem. Soc. 1950, 72, 2844–2847.

(39) Varadi, M. et al. AlphaFold Protein Structure Database: Massively Expanding the Structural Coverage of Protein-Sequence Space with High-Accuracy Models. Nucleic Acids Res 2022, 50, D439–D444.

(40) Hallgren, J.; Tsirigos, K. D.; Pedersen, M. D.; Armenteros, J. J. A.; Marcatili, P.; Nielsen, H.; Krogh, A.; Winther, O. DeepTMHMM Predicts Alpha and Beta Transmembrane Proteins Using Deep Neural Networks. bioRxiv 2022, 10.1101/2022.04.08.487609.

(41) Lambert, S. A.; Jolma, A.; Campitelli, L. F.; Das, P. K.; Yin, Y.; Albu, M.; Chen, X.; Taipale, J.; Hughes, T. R.; Weirauch, M. T. The Human Transcription Factors. Cell 2018, 172, 650–665.

(42) Larsen, F. B.; Voutsinos, V.; Jonsson, N.; Johansson, K. E.; Ethelberg, F. D.; Lindorff-Larsen, K.; Hartmann-Petersen, R. Comprehensive Degron Mapping in Human Transcription Factors. bioRxiv 2025, 10.1101/2025.05.16.654404.

(43) Heo, L.; Feig, M. One bead per residue can describe all-atom protein structures. Structure 2024, 32, 97–111.

(44) Grudinin, S.; Garkavenko, M.; Kazennov, A. Pepsi-SAXS: an adaptive method for rapid and accurate computation of small-angle X-ray scattering profiles. Biological Crystallography 2017, 73, 449–464.

